# Weak tension accelerates hybridization and dehybridization of short oligonucleotides

**DOI:** 10.1101/2022.04.19.488836

**Authors:** Derek J. Hart, Jiyoun Jeong, James C. Gumbart, Harold D. Kim

## Abstract

The hybridization and dehybridization of DNA subject to tension is relevant to fundamental genetic processes and to the design of DNA-based mechanobiology assays. While strong tension accelerates DNA melting and decelerates DNA annealing, the effects of tension weaker than 5 pN are less clear. In this study, we developed a DNA bow assay, which uses the bending rigidity of double-stranded DNA (dsDNA) to exert weak tension on a single-stranded DNA (ssDNA) target in the range of 2pN to 6pN. Combining this assay with single-molecule FRET, we measured the hybridization and dehybridization kinetics between a 15 nt ssDNA under tension and a 8-9 nt oligo, and found that both the hybridization and dehybridization rates monotonically increase with tension for various nucleotide sequences tested. These findings suggest that the nucleated duplex in its transition state is more extended than the pure dsDNA or ssDNA counterpart. Our simulations using the coarse-grained oxDNA2 model indicate that the increased extension of the transition state is due to exclusion interactions between unpaired ssDNA regions in close proximity to one another. This study highlights an example where the ideal worm-like chain models fail to explain the kinetic behavior of DNA in the low force regime.

## I. INTRODUCTION

DNA strand separation or unzipping followed by annealing or rezipping is commonplace in many fundamental genomic processes such as homologous recombination and R-loop formation [1-6]. Although genomic processes inside the cell are orchestrated by motor proteins or enzymes, they are thought to be aided by intrinsic dynamics of the underlying genomic DNA [7-11]. Therefore, thermally-induced separation of duplex DNA into single strands and its reverse reaction may play an important role in active genomic processes. For example, in both prokaryotic and eukaryotic genomes, origins of replication commonly feature a 10 bp to 100 bp DNA unwinding element, whose weak duplex stability determines origin function [12-14]. In CRISPR-Cas systems, melting is a rate-limiting step for Cas9 target selection, and has also been found to induce off-target binding and cleavage [15-17].

The melting probability of a duplex region depends not only on its sequence [18], but also on the local stress [18-22]. The genomic DNA *in vivo* is seldom in a relaxed state, but rather is subjected to various forms of stress: bending, twisting, and tension. Several DNA force spectroscopy experiments have carefully explored how melting is affected by a strong artificial tension [23-26], but the effect of weak tension (< 5pN), which is arguably more relevant to genomic processes *in vivo* or DNA-based systems *in vitro*, is less clear. Forces in this range can be exerted on a duplex region during active processes such as loop extrusion by SMC complexes [27, 28] and also by thermal fluctuations of flanking DNA segments [29]. Molecules involved in cell mechanotransduction also regularly experience forces at this scale [30]. Therefore, understanding the effect of weak tension on DNA hybridization/dehybridization can elucidate the physical regulation of genomic processes, and aid our design of DNA-based force sensors and actuators for the study of cell signaling mechanics [31-36] and the control of DNA nanostructures [37, 38].

In general, the force (*f*) dependence of two-state binding and unbinding kinetics can be modeled with a one-dimensional extension coordinate *x* as [39, 40]

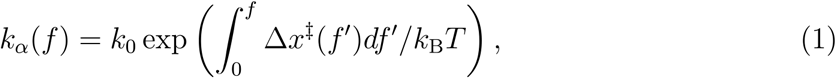

where *k*_*α*_ is the rate constant for binding (*α* = on) or unbinding (*α* = off), Δ *x* ^‡^ is the extension of the transition state *(x*^‡^) relative to the unbound *(x*_ub_) or bound state *(x*_b_), and *k*_B_*T* is the thermal energy. If the transition state is more extended than the bound state by a constant (Δ *x*^‡^ > 0), Equation 1 yields the well-known Bell’s formula [41]: *k*_off_ ∼ exp(*f*Δ*x*^‡^/*k*_B_*T*), which predicts that *k*_off_ monotonically increases with force. For DNA hybridization/dehybridization, the transition state is thought to be a nucleated duplex that contains both single-stranded DNA (ssDNA) and double-stranded DNA (dsDNA) [42-44]. According to the worm-like chain model, ssDNA, whose persistence length (*A*) is ∼ 1 nm, behaves like a flexible chain in the low force regime (*f* < *k*_B_*T*/*A* ∼ 5pN) [45]. It is thus conceivable that the transition state could be less extended than the pure dsDNA state in the low force regime (Figure 1A). Based on this idea, it was recently proposed that *k*_off_ (*f*) can decrease with force until *f* ∼ 5pN before increasing in the high force regime [40, 46, 47]. This counter-intuitive effect known as “roll-over” was predicted in a recent single-molecule fluorescence-tweezers experiment [48], but the limited data leaves the conclusion in question. Furthermore, how the extension of the nucleated duplex in the transition state compares to that of dsDNA in the bound state and pure ssDNA in the unbound state is not known.

**FIG. 1.**
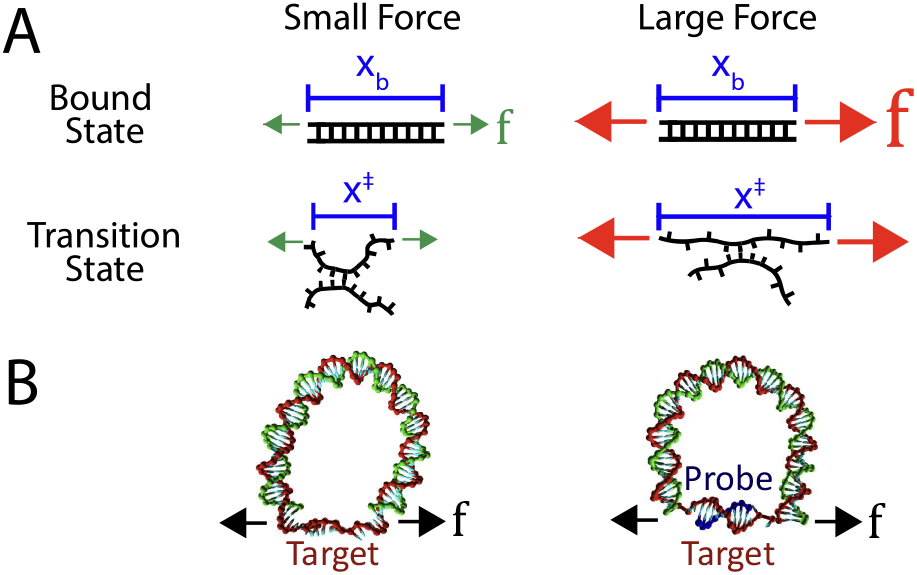
**(A)** A proposed model for how the extension of a duplex differs between small and large forces. The elasticity of the bound state is rigid, and therefore its extension *x*_*b*_ is mostly unaffected by force. On the other hand, the transition state may be more flexible, in which case its extension *x+* will be force-dependent. In this case, worm-like chain models predict that at large forces, *x*^‡^ > *x*_*b*_, whereas at small forces *x*^‡^ < *x*_*b*_*·* (B) DNA bow assay: a bent bow-like duplex of variable length exerts tension on a 15 nt ssDNA target (bowstring), extending the strand.

Here, we developed a DNA construct dubbed “DNA bow” (Figure 1B) to exert tension in the range between 2pN to 6pN on a short DNA oligo. The DNA bow is composed of a dsDNA segment (arc) of variable size (∼ 100 bp) and a short ssDNA target (bowstring); during experiment, a complementary ssDNA probe binds to and unbinds from this bow target. Combined with single-molecule FRET, DNA bows allow for high-throughput measurements of DNA hybridization and dehybridization kinetics in the low-force regime, using a conventional TIRF microscopy setup (Figure 2). Thus, this assay complements low-throughput, calibration-heavy tweezers [48, 49]. Using the DNA bow assay, we measured the hybridization and dehybridization rates of four DNA-DNA homoduplexes (lengths ranging from 8 bp to 9 bp) as well as their corresponding RNA-DNA heteroduplexes. Overall, the measured dehybridization (unbinding) rate monotonically increased with force with no clear sign of roll-over, and the measured hybridization (binding) rate also increased with force. In agreement with these experimental results, our simulations reveal that hybridization and dehybridization of short oligos transition through a maximally extended state, and as a result both processes are accelerated in the low force regime. We attribute the extension of the transition state to steric repulsion, which prevents the ssDNA overhangs of the nucleated duplex from coiling.

**FIG. 2.**
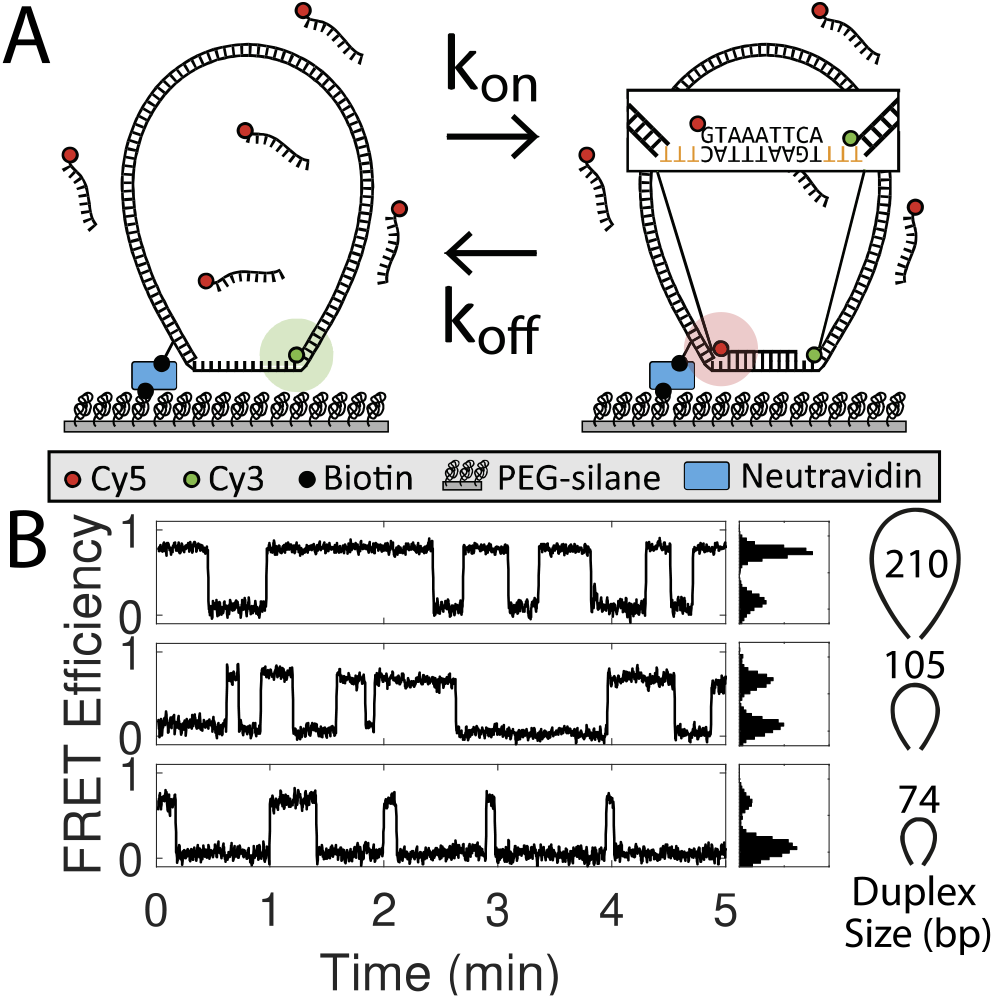
**(A)** Schematic of DNA bow assay FRET setup. Cy3-labeled DNA bows are immobilized on a PEGylated coverslip and excited by an evanescent wave of a 532-nm laser using TIRF microscopy. The inset highlights the ssDNA sequence (TGAAATTAC) targeted by the Cy5-labeled probe (GTAAATTCA). To avoid additional stacking interactions between the probe and the DNA bow, the 9 nt target segment was flanked by 3 nt ssDNA gaps (highlighted orange) in all construct designs. **(B)** Example FRET efficiency traces for three different dsDNA arc lengths (210, 105, 74 bp) exerting three separate forces (1.8, 3.8, 6.3 pN, respectively). FRET histograms are shown right. Binding and unbinding rates are extracted from the mean dwell times of low and high FRET states respectively.

## II. MATERIALS AND METHODS

### A. Preparing DNA bows

DNA bow molecules were constructed and labeled with a FRET donor (Cy3) and a biotin linker in 5 steps (Figure 3): (1) Template generation, (2) Modifier incorporation, (3) Circularization, (4) Nick generation, and (5) Strand exchange. Most notably, DNA bending protein HMG1 was used to facilitate intramolecular ligation of short DNA molecules [50].

**FIG. 3.**
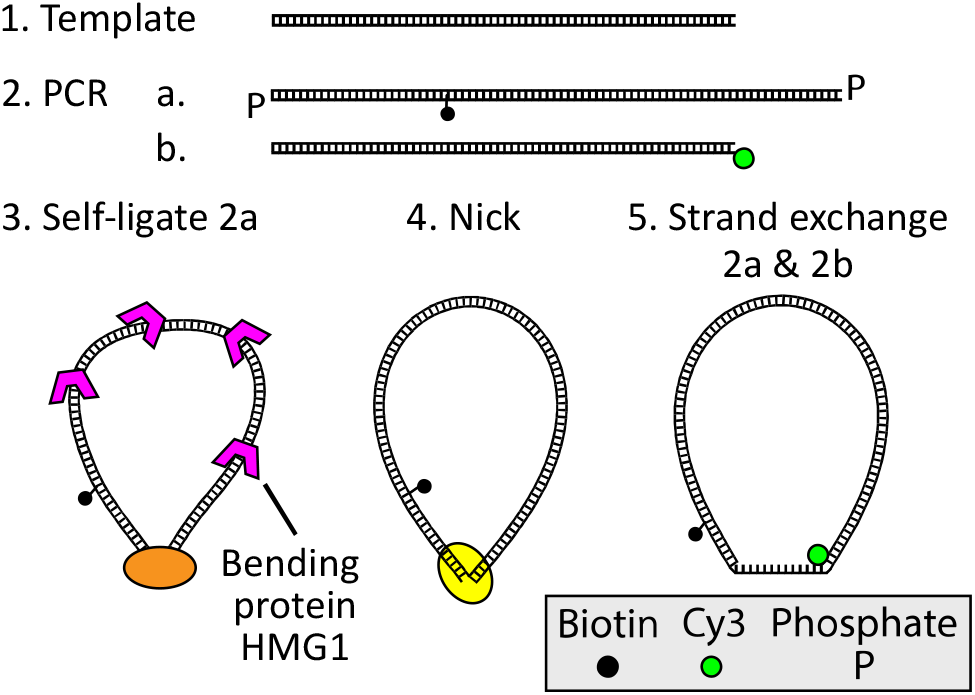
DNA bows were constructed in five steps. First, uniquely sized templates were generated with PCR from a common source. Using these templates, two sets of molecules (2a and 2b) were amplified with modified primers via PCR. DNA minicircles were then created from the phosphorylated 2a molecule set using protein-assisted DNA self-ligation. Afterward, DNA minicircles were purified and nicked on the unmodified strand. The final DNA bow constructs were finally constructed by exchanging the nicked strand of circularized 2a molecules with the Cy3-labeled 2b molecule.

In Step 1, polymerase chain reaction (PCR) was used to create a set of seven different DNA templates with lengths ranging from 74 bp to 252 bp (Supplementary Figure S3, Supplementary Table S1), using yeast genomic DNA as the source. The PCR primers were designed such that all seven templates shared adaptor sequences at their ends. In Step 2, using these templates, two additional PCR reactions were performed to create two sets of molecules, with each reaction using modified primers that anneal to the adaptor regions of the template. The first reaction produced a set of molecules with phosphorylated 5’ ends and an internal biotin-dT label for surface immobilization, as well as a 15 bp extension, consisting of a 9 bp target segment flanked on both sides by (dT)_3_ spacers. The second reaction produced donor-labeled (Cy3) molecules with a sequence identical to the original templates, which is 15 bp shorter than the first PCR product. All oligonucleotides were purchased from Eurofins MWC Operon and Integrated DNA Technology. All PCR products in the first and second steps were inspected by gel electrophoresis and extracted using a PCR clean-up kit. In Step 3, we circularized the phosphorylated molecules. To increase circularization efficiency, molecules were briefly incubated at 15 nM with 0.75 µM DNA bending protein HMG1 (Sigma Aldrich) in T4 ligase buffer for 10 minutes. Afterward, T4 ligase was added and the reaction volume was incubated overnight at 15 °C. The reaction was stopped via heat inactivation, after which T5 exonuclease was added to remove linear inter-molecular or nicked intra-molecular ligation products. Finally, Proteinase *K* was added to remove any protein leftovers. The remaining circular molecules were purified and concentrated using ethanol precipitation. In Step 4, the unmodified strand of our circular molecules was nicked using Nb.BbvCI in lx CutSmart buffer (NEB). After circularization and nicking, the resulting product was visualized and purified on a native polyacrylamide gel (6%, 29:1 acrylamide to bis-acrylamide in 0.5x TBE buffer), which appeared as a a single, isolated band as shown in Supplementary Figure S4. The bands were extracted using a simple “crush-and-soak” method, and then concentrated using the same ethanol precipitation method as before. In Step 5, a strand-exchange reaction was performed, replacing the nicked strand on each circular molecule with the corresponding donor-labeled linear strand. Circular molecules were mixed with the donor-labeled linear molecules at a 4:1 ratio, briefly heated to 95 °C, and gradually cooled down to 4 °C.

### B. DNA bow assay

Microscope slides with pre-drilled holes and coverslips were cleaned by sonicating in deionized water, drying in a vaccuum chamber, and 5-minute etching in a plasma chamber. The cleaned slides and coverslips were then passivated with PEG (polyethylene glycol) to minimize nonspecific binding. After PEGylation, the flow cell was assembled by joining the slide and the coverslip with double-sided tape and epoxy glue. The flow cell interior was incubated with NeutrAvidin followed by 50 µL of 40 pM DNA bow solution. Each measurement began after perfusing 20 nM of ssDNA probe solution into the flow chamber. The temperature of the flow chamber was maintained at 22 °C using an objective lens temperature controller. For each molecule, a high Cy3 signal (low FRET) indicates a DNA bow in the unbound state, while a high Cy5 signal (high FRET) indicates a DNA bow bound with the probe (Figure 2). Bound and unbound lifetimes of approximately ∼ 100 immobilized molecules were collected in each trial; 2-4 trials were performed for each bow size. All data was collected on an objective-based TIR microscope with an EMCCD camera (DU-897ECS0-BV, Andor). Frame times varied from 50 ms to 1000 ms, depending on the duplex sequence. The imaging buffer contained 100 mM NaCl, 100 mM Tris (8 pH), a triplet state quencher (1 mM Trolox), and the protocatechuic acid (PCA)/protocatechuate-3,4-dioxygenase (PCD) system [51]. Using this system, photobleaching was negligible at all donor excitation power settings and camera acquisition times used in our experiments.

### C. Data analysis

For each trial, time trajectories of FRET values were extracted from surface-immobilized molecules with in-house Matlab codes. Briefly, we calculated the FRET signal for each molecule from the background-subtracted intensities of the donor signal (*I*_*D*_) and the acceptor signal (*I*_*A*_) with *I*_*A*_/(*I*_*A*_ + *I*_*D*_). Next, we filtered FRET trajectories with a moving average, and used FRET signal thresholding to mark discrete transitions between the two FRET states. The dwell times in the bound (“on”) state (high-FRET state) and the unbound (“off’’) state (low-FRET state) were collected from each FRET trajectory. The binding rate (*k*_on_) and the unbinding rate (*k*_off_) were calculated from the mean dwell times (*τ*) using *k*_on_ = ([c] *τ*_off_)^−1^ and 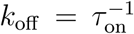, where [c] is the concentration of Cy5 labeled probes. Typically, ∼ 150 trajectories were used for each rate measurement.

### D. Estimating the tensile force exerted by a DNA bow

To estimate the tension exerted on the ssDNA bowstring, we treated the dsDNA arc as a worm-like chain. The force *f* exerted by a worm-like chain along its end-to-end direction at distance *x*_0_ can be calculated from the end-to-end distance (*x*) distribution *P*(*x*) of the chain according to

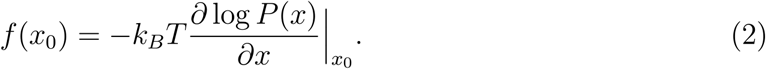

For *P*(*x*), we used an interpolated formula (Supplementary Equation S3), which is accurate for a wide range of bending stiffness values [52].

With our bow design, *x*_0_ also corresponds to the equilibrium extension of the ssDNA bowstring, and therefore its value will depend on both the bow size as well as whether the probe is bound to the complementary target segment. To find a realistic value of *x*_0_, we performed oxDNA2 simulations [53-55] for all possible combinations of bow size, target sequence, and probe state (bound or unbound). DNA bows bound to an RNA probe were not simulated; while oligomeric RNA-DNA duplexes have a slightly smaller helical rise [56], the overall effect that this difference would have on the force is negligible. Each MD simulation was run for *t* = 1.14 µs, using a time step of 15.2 fs. For each trajectory *n* = 7.5 × 10^4^ configurations were saved in 15.2 ps evenly spaced intervals. Using these saved configurations, we calculated the extension *x*, defined as the distance between the bases located at the terminal ends of the dsDNA bow and linked to the ssDNA target strand. The exact location of each terminal base was specified by its center of mass. Afterward, we calculated the mean extension 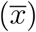 and standard deviation *σ*(*x*) for each molecule’s *x* distribution, to estimate *x*_0_ and its associated uncertainty respectively (Supplementary Table S2, Supplementary Figure S5). The tensile force 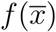 was then calculated using Equation 2, and the uncertainty in the tensile force was estimated with *σ*(*x*) by propagation of error, using 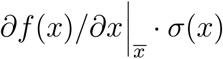. Additional details regarding WLC parameters and oxDNA2 simulations are provided in Supplementary Materials.

### E. Estimating hybridization and melting rates with FFS simulations

To determine hybridization and melting rates, we used a technique known as “forward flux sampling” (FFS) [57, 58]. This method ratchets the rare transition from an unbound state to a bound state, or vice versa, using a series of checkpoint interfaces which are each characterized by a unique order parameter value (e.g. minimum distance, number of bonds). By measuring the average flux Φ_*A*,0_ of a molecule in state A crossing the first interface *λ*_0_, and then measuring the probability *P*(*λ*_*Q*_|*λ*_*Q*_−1) of transitioning from *λ*_*Q*−1_ to *λ*_*Q*_ at each subsequent interface, it is possible to estimate the overall transition rate to state B (*k*_*AB*_), according to:

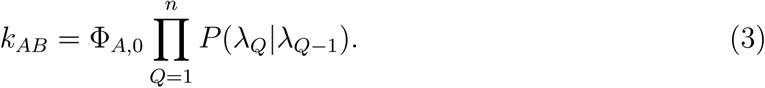

Using this technique with oxDNA2, the rate of the probe Pl-DNA binding to or unbinding from its corresponding target sequence T1 was calculated (Supplementary Table S1). Additional simulation details and parameter values are provided in Supplementary Materials.

### F. Observing the force-extension behavior of near-transition oligoduplexes

To measure the force-extension behavior of a partially melted oligoduplex near its binding or unbinding transition, we performed a series of MD simulations using the “mutual trap” external force tool provided with oxDNA2. Similar to the previous section, we simulated the target strand in four states: the “probe-bound” state (9bp), the ssDNA “probe-unbound” state (0 bp), a transition state with 1 bp remaining at the 3’ end of the duplex, and a transition state with 1 bp remaining at the center. In the transition state simulations, the remaining terminal or middle base pair interaction was strengthened 10-fold, while all other base pairing interactions were set to zero. For all simulations, the ends of the target strand were connected by a harmonic spring with stiffness *k* = 57.1pN nm^−1^ (1 simulation unit) and relaxed extension *x*_0_, such that the the tension *f* and extension *x* of the strand could easily be related using 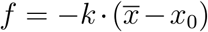. Similar to our DNA bow simulations, the extension *x* was defined as the distance between the center of mass of each terminal base on the target strand. For each state, we performed MD simulations for a small range of *x*_0_ values, such that the corresponding forces approximately spanned the force range of our DNA bows. For comparison, we plot the force-extension behavior of the target strand extended by a harmonic spring or a DNA bow in Supplementary Figure S6. Each simulation was performed for *t* = 1.52 µs using a time step of 15.2 fs. *n* = 10^5^ pairs of force and extension values were then calculated from configurations collected in 15.2 ps intervals evenly spaced across the MD trajectory.

## III. RESULTS AND DISCUSSION

Using the DNA bow assay, we measured the binding and unbinding rates of a short DNA or RNA (8-or 9-nt) oligo to a weakly pulled complementary target strand (15nt). The measured binding (*k*_on_) and unbinding (*k*_off_) rate constants thus reflect hybridization and dehybridization transitions of a short DNA homoduplex or DNA/RNA heteroduplex. Our DNA bow assay exploits the bending rigidity of dsDNA to generate small forces and is conceptually similar to the force clamp implemented with DNA origami [59] and a loop-based force transducer [60]. An identical DNA construct has also been used in other studies [61, 62]. Our DNA bow assay offers unique advantages over other single-molecule force assays such as optical and magnetic tweezers in that (1) force measurements can be performed on many molecules in parallel, and (2) no calibration of force vs. extension is required for each molecule since the force is generated by chemically identical DNA molecules, not through beads of variable properties. We created 21 DNA bows in total, including 7 different dsDNA lengths (74, 84, 105, 126, 158, 210 and 252 bp) for the elastic arc segment and 3 unique sequences for the complementary segment of the ssDNA target. The DNA bow was further designed such that only the desired gapped DNA circle can generate the FRET signal from the surface upon probe binding (Supplementary Figure S7). While sharp bending is known to disrupt the helical structure of circular DNA by generating “kinks”, these deformations do not appear in circles larger than 84 bp [63]. Therefore, we predict that kinking is negligible even for our smallest DNA bow size, which includes a flexible 15 bp ssDNA segment in addition to its 74 bp dsDNA arc. Given this, the force generated by each DNA bow was calculated by treating the DNA arc as a simple worm-like chain. Using this assumption, the range of forces exerted on the target strand is calculated to be 1.70 pN to 6.34 pN in the unbound state and 1.6 pN to 6.25 pN in the bound state.

### A. Binding and unbinding rates vs. force

In Figure 4, we present the force dependence of *k*_on_ (A) and *k*_off_ (B) for 4 DNA-DNA duplexes (left column) and 4 RNA-DNA duplexes (right column). The scale of y-axis is set as logarithmic to aid comparison to Equation 1. Each RNA sequence is identical to a corresponding DNA sequence, except for T to U substitution. As shown in Figure 4A, *k*_on_ tends to increase with force over the measured force range. The relative increase in *k*_on_ is sequence-dependent: the increase is relatively large for AGGACTTGT but small for GTAAATTCA. The relative increase or dynamic range is quantified by taking the ratio of the rate at the highest force to that at the lowest force (Figure 4C). This sequence-dependence was also observed in RNA-DNA duplexes, with each heteroduplex approximately matching the behavior of its corresponding homoduplex. However, these differences in relative increase mostly disappear above 3pN, and *k*_on_ appears to reach a plateau above 6pN. We note that the comparison of the second-order rate constant *k*_on_ across different sequences is not accurate because of the inaccuracy in the estimated concentration of each probe.

**FIG. 4.**
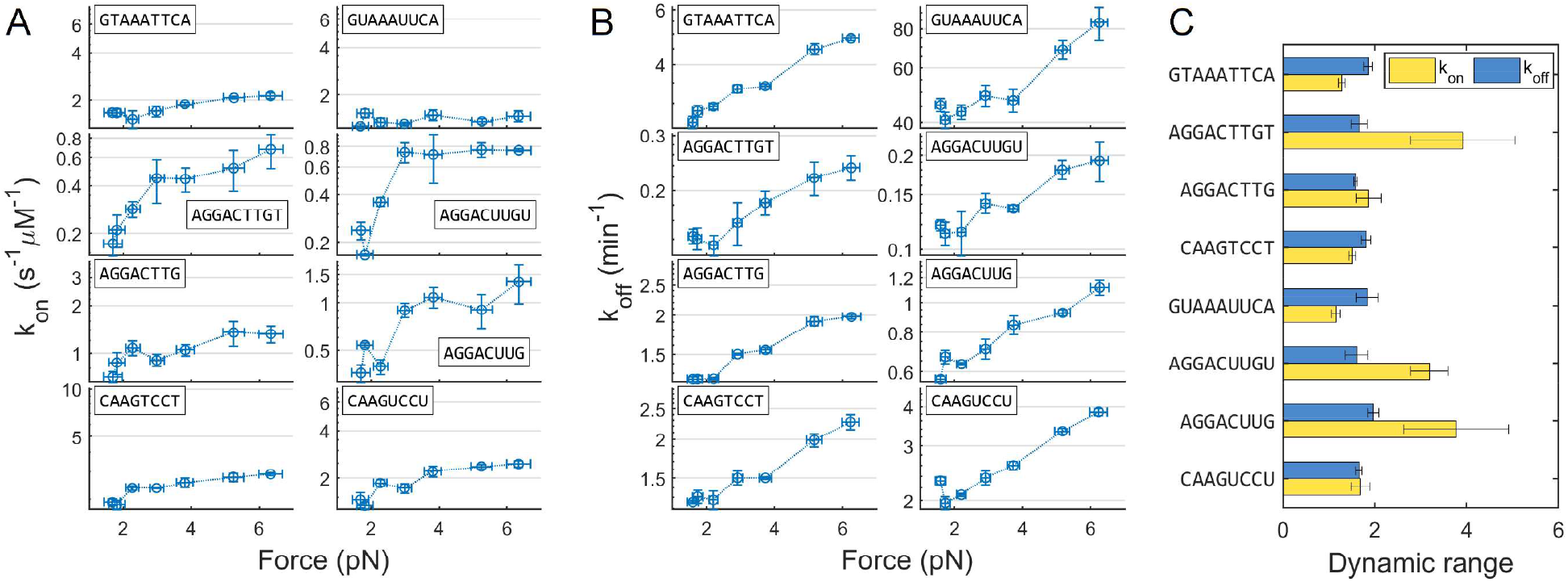
**(A)** Binding rate vs. force. The plots on the left (right) column are for DNA (RNA) probes. The y-axis is on a logarithmic scale over the same 5.8-fold change for all probe sequences. **(B)** Unbinding rate vs. force. The plots on the left (right) column are for DNA (RNA) probes. The y-axis is on a logarithmic scale over the same 2.4-fold change for all probe sequences. Vertical error bars for binding and unbinding rates represent the standard error of the mean; horizontal error bars were calculated using 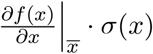, where 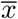 and *σ*(*x*) are the mean and standard deviation of the bow’s end-to-end distance distribution. **(C)** The dynamic range of all measured rates. The dynamic range was obtained by dividing the rate at the highest force by that at the lowest force; the associated error was calculated by propagating the uncertainty in the underlying rates.

The most significant result from Figure 4A is that *k*_on_, the binding rate of the probe to its complementary target, becomes faster, not slower, as the tension in the target strand increases. This result stands in contrast to previous rates observed at higher forces, such as those observed for DNA hairpin folding [64-66]. By differentiating the logarithm of Equation 1 with respect to force, we can relate the slope of curves in Figure 4A to Δ*x*^‡^, which is the extension of the transition state (*x*^‡^) relative to the unbound state (*x*_ub_):

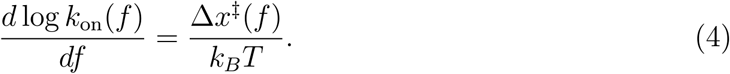

The overall non-negative slope in Figure 4A indicates that the transition state for hybridization is more extended than the unbound state (*x*^*‡*^ > *x*_ub_) in the range of 2 pN to 6 pN.

The force dependence of *k*_off_ is shown in Figure 4B. Compared to *k*_on_, the dynamic range for *k*_off_ is somewhat uniform at 2-fold across all DNA-DNA and DNA-RNA duplexes (Figure 4C). The apparent slope is mostly positive except between a few points below 2pN, which implies that the roll-over effect or catch-to-slip transition is negligible. Similar to Equation 4, the slope of curves in Figure 4B is proportional to Δ*x*^‡^ for dehybridization, which is the extension of the transition state (*x*^‡^) relative to the bound state (*x*_b_)-From this, we conclude that the transition state for dehybridization is more extended than the bound state (*x*^‡^ > *x*_b_) in the range of 2pN to 6pN.

Since the first-order rate constant *k*_off_ is concentration-independent, it can be compared across different sequences. When compared at the same force, *k*_off_ was in the order of GTAAATTCA > AGGACTTG = CAAGTCCT > AGGACTTGT from fastest to slowest. When a single nucleotide was removed from the 3’ end of AGGACTTGT, *k*_off_ increased as expected from the weaker base pairing interaction. Between AGGACTTG and its reverse complement CAAGTCCT, *k*_off_ remains the same, which implies that for a DNA-DNA homoduplex, *k*_off_ is similar regardless of which strand is subject to tension. *k*_off_ for RNA-DNA duplexes (Figure 4A, right) similarly showed a strong sequence-dependence, in the order of GUAAAUUCA>CAAGUCCU>AGGACUUG>AGGACUUGU. In two cases (AGGACUUGU and AGGACUUG), RNA-DNA heteroduplex was longer-lived than its homoduplex counterpart, but in the other two (GUAAAUUCA and CAAGUCCU), DNA-DNA homoduplex was longer-lived.

### B. Thermodynamic stability

From the individual rate constants, we can calculate the standard free energy difference (Δ*G*°) between the bound and unbound states according to

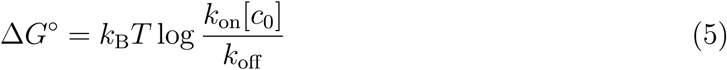

where [*c*_0_] is 1 M. In this definition, ΔG^0^ is more positive for a more stable duplex. In Supplementary Figure S8, we compare ΔG^0^ calculated using 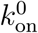 and 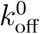 with 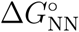 estimated using a nearest-neighbor (NN) thermodynamic model [67]. Most sequences are significantly more stable than the model predicts, showing at least a 2 *k*_*B*_*T* difference. This increased stability can be attributed to two major factors. First, the terminal bases of the duplex will stack with the adjacent unpaired bases in the gaps, which has been shown to provide ∼ 1 *k*_*B*_*T* per end interaction in 8 bp DNA duplexes [68]. Dangling nucleotides beyond these adjacent bases have also been shown to stabilize the duplex [69, 70], albeit to a lesser degree [71]. Second, the DNA and RNA probes used in this experiment were labeled with a Cy5 dye on the 5’ end, which will also stabilize short DNA duplexes by 2 *k*_*B*_*T* [72]. The stabilizing effects of dangling-base interactions and 5’ dye labeling are additive [72]. When accounting for these two factors, we find that the nearest-neighbor model prediction matches Δ*G*° more closely.

In Supplementary Figure S9, the free energy difference Δ*G*° is plotted against force. Because both rates change in the same direction in response to force, the force dependence of Δ*G*° is somewhat dampened. Except for AGGACTTGT and its RNA counterpart, Δ*G*° changes little, albeit with some scatter. In comparison, the force-dependence of Δ*G*° of AGGACTTGT and AGGACUUGU shows a monotonic increase up to 3pN and afterward plateaus, varying by less than 0.5 *k*_B_*T*.

### C. oxDNA2 simulations of binding and unbinding trajectories

To gain molecular insights into binding and unbinding transitions, we performed coarsegrained simulations of both reactions using oxDNA2. For both simulations, the probe Pl-DNA was simulated together with its corresponding target sequence T1 (Supplementary Table S1). The end-to-end extension of the target strand was held fixed using the harmonic trap tool provided with oxDNA2, while the probe was allowed to diffuse freely. The target strand was held at 5.5 nm and 5.1 nm for unbinding and binding reactions, respectively. These values were determined from the average target strand extensions of the largest DNA bow in the bound and unbound states. Because binding and unbinding events are relatively rare, we implemented forward flux sampling, which separates rare events into computationally feasible intervals [57, 58]. Each interval was demarcated by two interfaces, where each interface was defined using a relevant order parameter (Supplementary Tables S4 and S3). Using Equation 3, we calculated an average *k*_on_ value of 2.3 s^−1^ µM^−1^ (where the probe concentration was estimated using the [10.2 nm]^3^ simulation box volume) and an average *k*_off_ value of 9.9 min^−1^. A direct comparison between the calculated rates and the measured rates is not accurate considering that coarse-graining is known to speed up dynamical timescales by smoothing energy landscapes and neglecting hydrodynamic effects [73-75]. Nonetheless, the reaction paths should shed light on the nature of the transition states.

The initial unbinding rate of the first-melted base, as well as the melting probability of each subsequent base, are individually tabulated in Table S5. An additional step is also included which calculates the probability that the strands exceed a minimum distance *d* = 0.85nm after all bases have melted (Table S3). Throughout unbinding, the probability of melting each base is relatively small, ranging from 0.03 to 0.06. Notably, the chance that oligos separate from one another remains low even after all bases have melted: according to the final strand separation step, a newly-melted duplex is expected to re-form a base pair with *p* ∼ 0.9 probability. This fast reassociation between short oligos is similar to that recently observed between DNA and the lac repressor [76]. This result also suggests that the unbinding transition state happens after all base pairs have already melted. Therefore, the apparent transition state for a FRET-based dissociation event should occur after the oligos are physically well separated by some distance.

We also measured the average base-pairing and stacking potentials of the complementary target segment for all unbinding steps (Figure 5A). As the unbinding reaction progressed, both pairing and stacking potentials weakened in a symmetric fashion; interactions on the edges of the duplex were more likely to be broken than interactions nearest to the center. Stacking interactions were largely unaffected during the first 3 melting steps (> 6bp), and remain relatively high up to the final base melting step (0 bp). During the last few steps (3bp to 0 bp), the remaining interactions of GTAAATTCA are skewed towards one side in an apparent symmetry breaking, centered on the relatively strong CT dinucleotide pair. The discrete change in the stacking potential after strand separation (*d* > 0.85 nm) suggests that the oligos maintain some residual helical stacking immediately after duplex melting, which may enable the fast reassociation previously discussed.

**FIG. 5.**
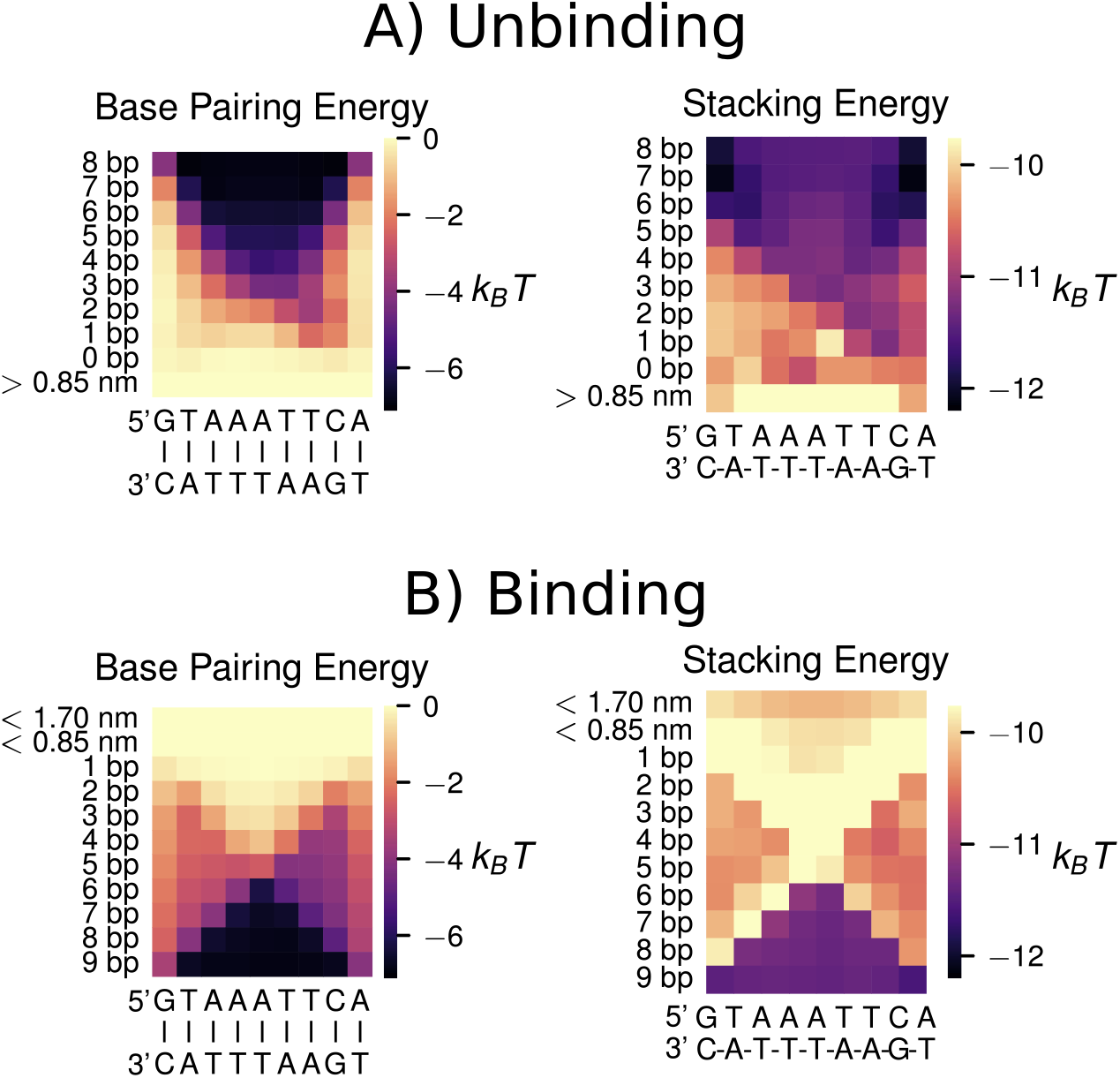
Average pairing and stacking potentials for the complementary portion of the target strand at each interface of **(A)** binding and **(B)** unbinding FFS simulations. Colorbar ticks are specified in *k*_*B*_*T* units, where T = 22 °C.

Similar to unbinding, the individual steps of binding are also enumerated in Table S5. They include two strand approach steps (where interfaces are defined by inter-strand separation going below a minimum distance threshold) as well as nine base pairing steps (Table S4). The slowest step in the binding process was the formation of the first base pair (starting at minimum distance *d* = 0.85nm between matching bases). The success probability of this step was less than 5 × 10^−3^, an order of magnitude lower than in any other step. After this step, however, the probability of additional base pairing increases rapidly, in stark contrast to the low probabilities seen throughout the unbinding reaction. The likelihood of full duplex formation approaches one after only four bases have formed, consistent with the zipping model for DNA binding [77, 78].

As before, we measured the pairing and stacking potentials of both sequences as binding progressed (Figure 5B). In stark contrast to unbinding, all pairing potentials first strengthen at either end of the target strand and afterwards “zip” in a linear fashion. While both pathways were common, we observed a slight 5’ to 3’ preference for the target strand zipping direction. Stacking potentials increased in strength in a similar fashion; however, these potentials temporarily weaken just before pairing occurs. This result suggests that local unstacking may promote duplex nucleation by granting more orientational freedom to bases.

Comparing these reactions to unbinding, we find that hybridization is not a simple reversal of melting (Supplementary Figure S10). Melting adopts a “fray and peel” pathway, where unbinding begins at the duplex termini and slowly proceeds base by base toward the duplex center. By contrast, hybridization follows a “dock and zip” pathway, where the strands anneal with high probability after the rare formation of a toehold at a strand terminus. Together, these differences demonstrate that binding and unbinding take different reaction pathways and do not share the same transition barrier.

### D. The physical nature of the transition state(s)

Our DNA-bow experiments show that in the force range of 2 pN to 6 pN, both the binding and unbinding rates increase with force. The fact that weak force increases the accessibility of the transition state implies that the transition state is more extended than the two observable states, bound and unbound. At the same time, our kinetics simulations show that hybridization and dehybridization do not share the same transition state. Hybridization more likely occurs through the formation of a few terminal base pairs (end-paired transition state) while dehybridization occurs after the breakage of the last remaining base pair in the center (middle-paired transition state). To rationalize our experimental results with extension *x* as the sole reaction coordinate, we obtained the force-extension curves of bound, unbound, and two different transition states (Figure 6A) from oxDNA2 simulations. Since the transition state is too transient to be analyzed in a normal dynamics simulation, we stalled the system near this state by turning off all base pairing interaction except in one central or terminal base pair, whose pairing interaction was strengthened 10-fold. As shown in Figure 6B, we find that the transition state for hybridization (end-paired) is more extended than the unbound state (ssDNA), and the transition state for dehybridization (middle-paired) is more extended than the bound state (dsDNA) over the entire force-range of our experimental assay. Hence, our simulation results are consistent with the measured force dependence of both *k*_on_ and *k*_off_.

**FIG. 6.**
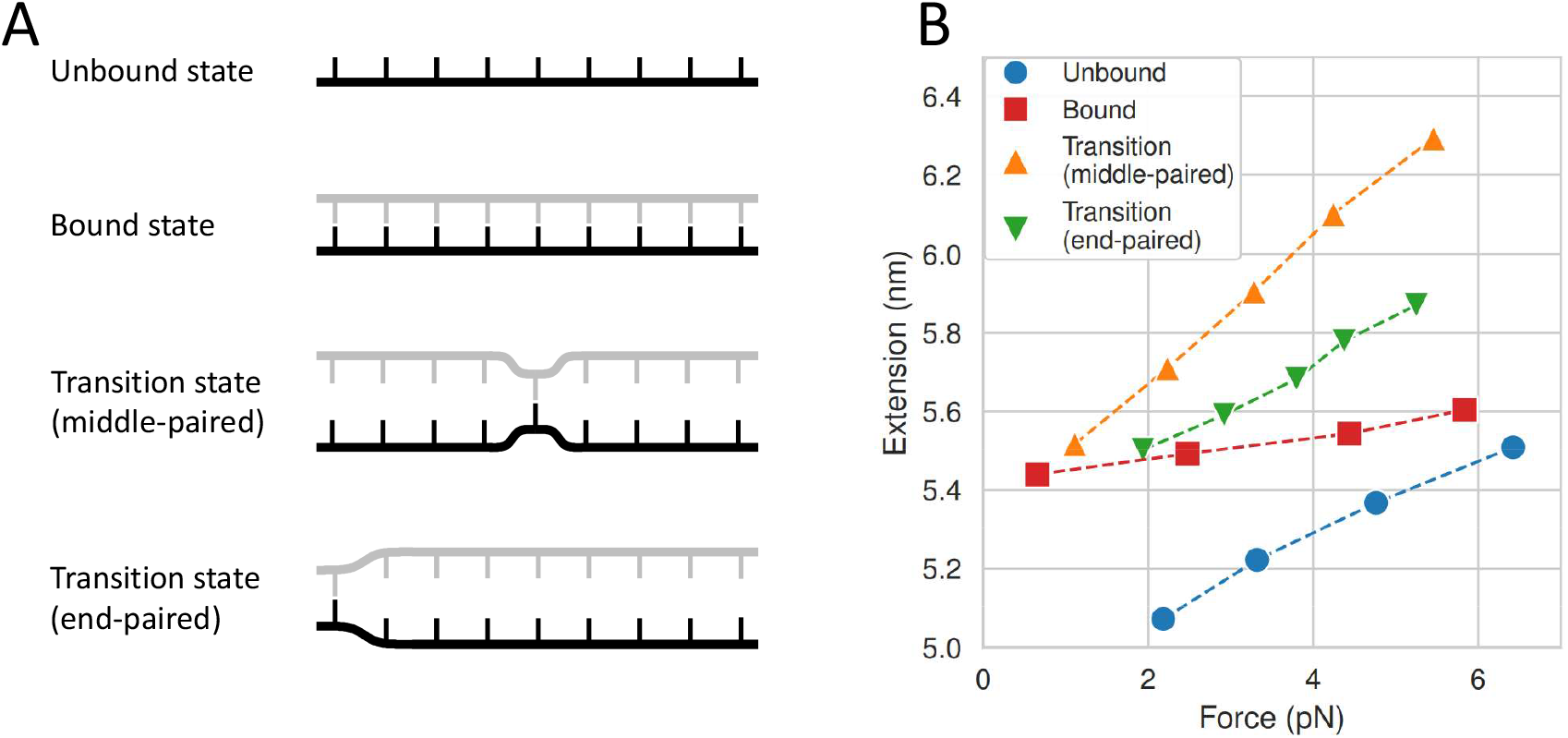
**(A)** Schematic of target strand (black) in four unique states: the unbound state, the bound state, the transition state with one base pair in the middle, and the transition state with one terminal base pair. **(B)** Force-extension curves of target strand in each state. For each filled marker, a unique MD simulation of the target segment was performed (*t* = 75.75 ns, *dt* = 15.2 fs). Afterward, the extension value was estimated by calculating the average distance 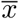 between the center-of-masses of the terminal bases on the target strand, and force values were calculated using 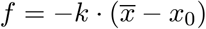 Dotted lines represent a linear regression of force-extension values observed for state. For comparison, the average extension values and corresponding force values of all bow sizes containing the same target sequence are also shown, in both its bound and unbound states.

At first sight, it is not obvious why the transition state, which is a mixed state of ss-DNA and dsDNA, is more extended than the bound and unbound state, which are pure dsDNA and ssDNA, respectively. We speculate that ssDNA strands confined to close proximity prevent each other from adopting randomly coiled conformations. To the same effect, randomly coiled conformations of ssDNA strands are not compatible to form a nucleated duplex. Therefore, ssDNA regions in the transition state happen to be more extended than in isolation.

We find that the higher extension of DNA in its transition state is likely created by exclusion interactions between the target and probe strands. When comparing the unbound state to the binding transition state, bases nearby the end pair location were on average much further from bases on the opposite side of the strand, suggesting that the presence of the probe in the binding transition state blocks folded conformations. This may explain why weak tension increases the accessibility of the transition state, which extends ssDNA without overstretching, bringing the target strand closer to its “unfolded” transition state. For the unbinding transition, melted bases at one end of the complementary segment were much further from bases at the opposite end. This increased distance between the ssDNA overhangs occurs in spite of their increased flexibility, presumably due to exclusion interactions that occur between the target and the probe.

### E. Comments on roll-over or catch-to-slip transition

The roll-over effect was postulated based on the idea that the transition state is a hybrid of ssDNA and dsDNA and that each obeys the force-extension formula for an ideal WLC [40, 47] or a unique WLC with its own characteristics [48]. The interpolation formula used in these models, however, is not valid for short chains [79, 80]. For example, the formula predicts that the average end-to-end distance of a 10-nt ssDNA or dsDNA is zero, which is obviously incorrect. A more accurate formula derived for short chains [81, 82] places the crossover force at ∼ 1.8pN (Supplementary Note), which borders the force limit of our DNA bow assay. However, even this formula cannot accurately describe the force-extension behavior of a nucleated duplex, whose unpaired regions are unavoidably influenced by exclusion interactions. Instead, we used oxDNA2 simulations to directly obtain the force-extension curves of the short ssDNA, dsDNA, and transition state. As shown in Figure 6B, the transition state is more extended than either ssDNA or dsDNA across the entire force range of our experimental assay. However, our study does not completely eliminate the possibility of a roll-over. First, our DNA bow assay cannot probe forces lower than 1.5 pN. In this range, we find that the extension of the transition state can become shorter than that of the bound state (Figure 6). Second, the roll-over effect is predicted to be more pronounced for longer oligos [47]. A more thorough test of this model thus requires measuring the dehybridization rate of oligos longer than 10 nt, which is extremely slow (∼ hr^−1^). Therefore, the roll-over effect, if any, would only exist on a time scale too slow to bear any physiological or practical significance beyond the theoretical realm.

## IV. CONCLUSION

DNA often experiences tension through passive or active mechanisms. In the presence of 5pN of force, DNA polymer models predict that dsDNA and ssDNA have a similar extension, which can lead to a nontrivial force dependence of hybridization and dehybridization rates. Previous force spectroscopy techniques, however, are not suitable for investigating this force dependence due to limited throughput. In this study, we developed a DNA bow assay, which can exert 2 pN to 6 pN of tension on a ssDNA target and report on its hybridization and dehybridization via smFRET. In this force range, we found that both the hybridization and dehybridization rates increase with force, which indicates that the transition state has a longer extension than its ssDNA and dsDNA counterparts. Coarse-grained simulations reveal that hybridization and dehybridization proceed through different transition states with a single base pair formed near the end or the middle. Consistent with the experimental results, simulations also show that these two transition states are indeed more extended than their respective initial states due to exclusion interactions that preclude ssDNA overhangs from adopting random coil configurations. Our study underscores the importance of investigating DNA-based reaction kinetics in the low force regime, which are not predictable by canonical polymer models of DNA.

## V. DATA AVAILABILITY

All data presented in this manuscript can be made available upon request from the corresponding author.

## VI. ACKNOWLEDGEMENTS

The authors thank the members of the Kim laboratory for useful discussions. Computational resources were provided by the Partnership for an Advanced Computing Envrironment (PACE) at the Georgia Institute of Technology.

