## Supplementary Material for "Weak tension accelerates hybridization and dehybridization of short oligonucleotides"

### Supplementary Note: Force-extension curves of worm-like DNA

In the limit of $L \gg A$, Marko and Siggia derived an interpolation formula (MS formula) for the relationship between force ($f$) and extension ($x$) of a worm-like chain (WLC) [79]:

$$\frac{fA}{k_{\mathrm{B}}T} = \frac{x}{L} + \frac{1}{4\left(1 - x/L\right)^2} - \frac{1}{4} \tag{S1}$$

where $A$ is the persistence length, and $L$ is the contour length. It is convenient to define the contour length per nucleotide, $b = L/N$. The accuracy of this formula can be increased with additional terms [80]. Whitley et al. [48] and Guo et al. [46] modeled ssDNA and dsDNA as WLCs and also attempted modeling the transition state as a chimeric DNA of ssDNA and dsDNA or a WLC with its own unique $A$ and $L$. Using 53 nm and 0.34 nm for $A$ and $b$ of dsDNA, and 1.32 nm and 0.6 nm for $A$ and $b$ of ssDNA in Equation S1 and inverting it, we can obtain $x$ as a function of $f$ (top, Supplementary Figure S1). The extensions of

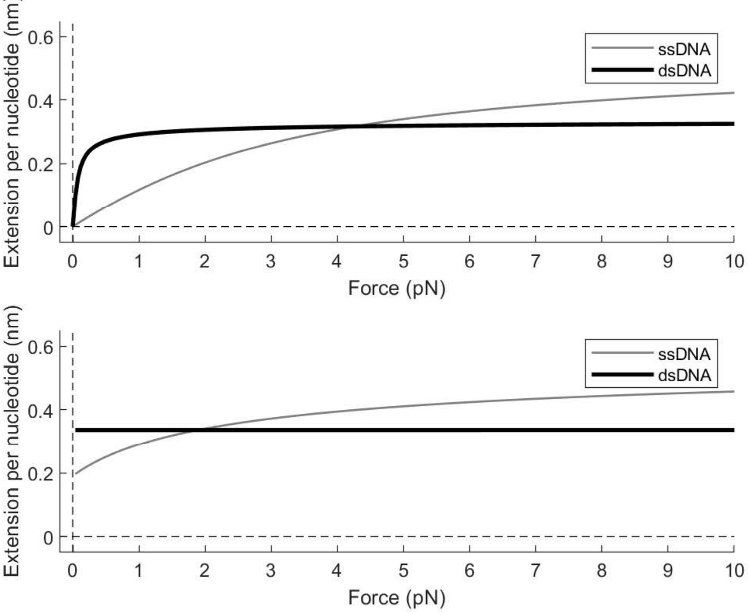


Supplementary Figure S1.

ssDNA and dsDNA are predicted to cross over at $f \approx 4.3$. A different formula that is more correct for short WLC is derived by Keller et al. [81] and Hori et al. [82]. In this formula,

$x$ is expressed as a function of $f$:

$$x = L - \frac{k_\mathrm{B}T}{2f}\left(L\sqrt{\frac{f}{Ak_\mathrm{B}T}}\coth\left(L\sqrt{\frac{f}{Ak_\mathrm{B}T}}\right) - 1\right). \tag{S2}$$

Force-extension curves of ssDNA and dsDNA obtained from this formula are shown at the bottom of Supplementary Figure S1. The crossover force is $\sim 1.8$ pN, markedly lower than predicted by the MS formula.

### A. The role of stacking in hybridization

Previous studies have hypothesized that weak DNA tension promotes binding by ordering ssDNA into "prehelical" structures [83, 84]. However, when comparing the average stacking interactions of the ssDNA target strand for all DNA bow sizes, we observe that stacking interactions are weakest in our smallest DNA bows, for which our experimental results show higher hybridization rates (Supplementary Figure S2). Moreover, our kinetics simulation

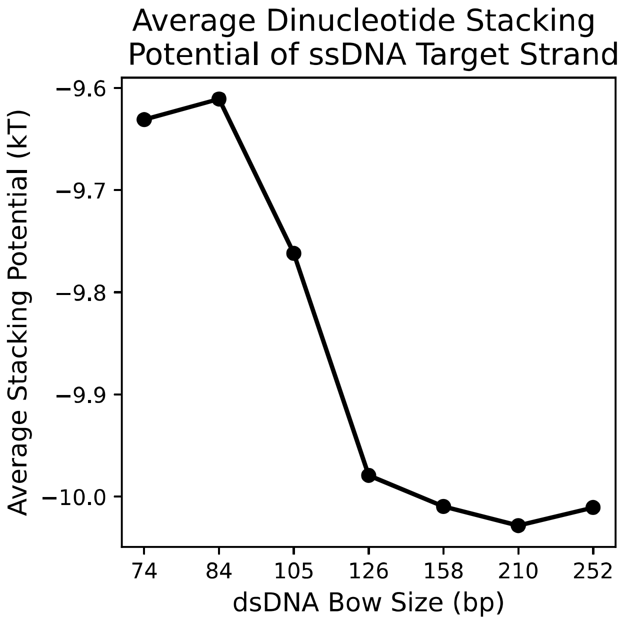


Supplementary Figure S2. Average dinucleotide stacking potential of the ssDNA target sequence T2 for all DNA bow sizes. Energy values for each bow size are averaged over all saved configurations $7.5\times10^4$ and over all dinucleotide pairs in the target region. Note that stacking interactions between the terminal base pair of the dsDNA bow and the adjacent unpaired base in the ssDNA target are included in the average.

indicates that stacking itself may have a negative impact on initial pairing between two complementary strands (Figure 5). These two results are counterintuitive given that stacking stabilizes dsDNA. However, for bases on opposite strands to pair, they need rotational freedom, which would be restricted if base stacking is present. Therefore, base stacking seems to play conflicting roles in both hindering base pair formation prior to strand docking as well as stabilizing base pairs once they are formed.

**Supplementary Information: oxDNA2 simulation parameters**

For all oxDNA2 simulations, the buffer conditions were specified to be identical to our experiments (100 mM salt concentration and 22 °C). To prevent non-representative initial states, all simulations were equilibrated for 50000 time steps before configurations were saved into output trajectories [73]. All simulations used an Andersen-like thermostat [85], where the molecular system was propagated according to Newton's equations for $N_{\mathrm{Newt}}$ time steps using Verlet integration; afterward, the system was assigned new linear and angular velocities drawn from a Maxwell-Boltzmann distribution such that the resulting diffusion coefficient was equal to a specified value $D$. For all DNA bow simulations and force-extension simulations, $N_{\mathrm{Newt}} = 103$ and $D = 2.5$; for all FFS simulations, $N_{\mathrm{Newt}} = 51$ and $D = 1.25$.

**Supplementary Method: Estimating the end-to-end distance radial probability distribution of DNA bows**

To estimate the tensile force $f$ exerted by each DNA bow size (Equation 2), we used the following interpolation formula to estimate the radial probability distribution $P(x)$ of a wormlike chain

$$\begin{aligned} P(x') = 4\pi x'^{\,2} \cdot J_{SYD} \cdot \left(\frac{1-cx'^2}{1-x'^2}\right)^{5/2} \exp\left(\frac{\sum_{i=-1}^{0}\sum_{j=1}^{3} c_{i,j}\kappa^i x'^{2j}}{1-x'^2}\right) \\ \times \exp\left(\frac{-d\kappa ab(1+b)x'^2}{1-b^2x'^2}\right) I_0\left(-\frac{d\kappa ab(1+b)x'^2}{1-b^2x'^2}\right), \end{aligned} \tag{S3}$$

where

$$a = 14.054, \quad b = 0.473, \quad (c_{i,j})_{i,j} = \begin{pmatrix} -3/4 & 23/64 & -7/64 \\ -1/2 & 17/16 & -9/16 \end{pmatrix}.$$

This formula accurately models $P(x)$ for a large range of stiffness values ($\kappa = A/L$, where $A$ and $L$ are the persistence and contour lengths of the dsDNA elastic arc, respectively) as well as a wide range of normalized end-to-end distance values [52] ($x' = x/L$). For this calculation, we assumed the values $A = 53\,\mathrm{nm}$ and $b = 0.34\,\mathrm{nm}$, where $b$ is the contour length per nucleotide $b = L/N$ Therefore, Equation S3 can be used with Equation 2 to estimate the force exerted by all bow sizes, whose stiffnesses range from $\kappa = 0.6$ to $\kappa$=2.1, and whose end-to-end distance values range from $x = 0.06$ to $x = 0.21$.

**Supplementary Method: Simulating unbinding and binding reactions with forward flux sampling (FFS)**

For all FFS simulations, the center of mass of each of the terminal bases on the 17 nt target molecule T1 were separated by a fixed extension value $x$ using two strong harmonic traps with force constant k = 570.9 pN/nm (10 simulation units in oxDNA). $x$ was fixed at 5.5 nm for unbinding reactions and 5.1 nm for binding reactions, which are equivalent to the average extension values observed for our largest DNA bow in its bound and unbound state respectively (Tables S2). All FFS simulations were performed using a 9.1 fs time step.

The unbinding FFS simulation was separated into 9 interfaces (Supplementary Table S3. Each interface $\lambda_Q$ corresponded to a change in the number of remaining base pairs, decreasing from 8 remaining base pairs to 0 base pairs. Base pairing was defined as when any two complementary bases had a hydrogen bond potential energy less than -0.1 simulation units ($-0.6$ kcal/mol, or about 1 $k_BT$ at 22 °C). The initial flux was calculated by running a brute force trajectory of the molecule in state A (9 bp) and observing the rate of forward crossings across the first interface $\lambda_0$ (8 bp) according to

$$\Phi_{A,0} = \frac{N_0}{T}, \tag{S4}$$

where $N_0$ is the number of crossings and T is the total time duration of trajectories where A was more recently visited than B. Using the configurations of successful crossings saved during initial flux simulation, the transition probability $P(\lambda_1|\lambda_0)$ of melting the next base pair (8 bp to 7 bp) was calculated according to

$$P(\lambda_1|\lambda_0) = \frac{N_1}{M_0}, \tag{S5}$$

where $M_0$ is the number of trial trajectories started at $\lambda_0$ and $N_1$ is the number of trajectories that successfully reach $\lambda_1$. Note that trajectories are halted and marked as failures upon reaching state A. Each trial trajectory is started from a configuration randomly selected from the $N_0$ configurations saved during the initial flux simulation. In the following steps, we implemented a variant of FFS known as "pruning", which eliminates a large fraction of backward-moving trajectories and re-weights the surviving trials [86]. In these steps, trajectories that revert backward to interface $\lambda_{Q-2}$ after starting at $\lambda_{Q-1}$ were pruned with probability $p = 0.75$. To correct for this, the transition probability $P(\lambda_Q|\lambda_{Q-1})$ of melting additional base pairs was calculated according to

$$P(\lambda_Q|\lambda_Q - 1) = \frac{N_Q - N_Q^* + N_Q^*/(1-p)}{M_{Q-1}}, \tag{S6}$$

where $M_{Q-1}$ is the number of trial trajectories started at $\lambda_{Q-1}$, $N_Q$ is the total number of trajectories that successfully reach $\lambda_Q$, and $N_Q^*$ is the number of trajectories that revert to $Q-2$, survive pruning, and ultimately reach $\lambda_Q$. The initial flux of crossing $\lambda_0$, as well as the probabilities of crossing successive interfaces, are tabulated in S5.

The binding FFS simulation was separated into 11 interfaces. Similar to unbinding, the initial flux as well as the subsequent melting probabilities were calculated using Equations S4–S5. As before, pruning was implemented for interfaces after $\lambda_0$. Interfaces for the first two "strand approach" steps were defined with a distance order parameter $d$, where $d$ was defined as the minimum separation between any two complementary bases on the probe and target segment. Similar to unbinding, interfaces for the remaining 9 steps corresponded to a change in the number of paired bases $n$ in the partial duplex, starting at 1 bp and ending at 9 bp (Supplementary Table S4). Similar to unbinding, base pairs were defined as when any two complementary bases had a hydrogen bond potential energy of less than -0.1 simulation units ($-0.6$ kcal/mol). The initial flux of crossing $\lambda_0$, as well as the probabilities of crossing each successive interface ($\lambda_Q$), are tabulated in S5.

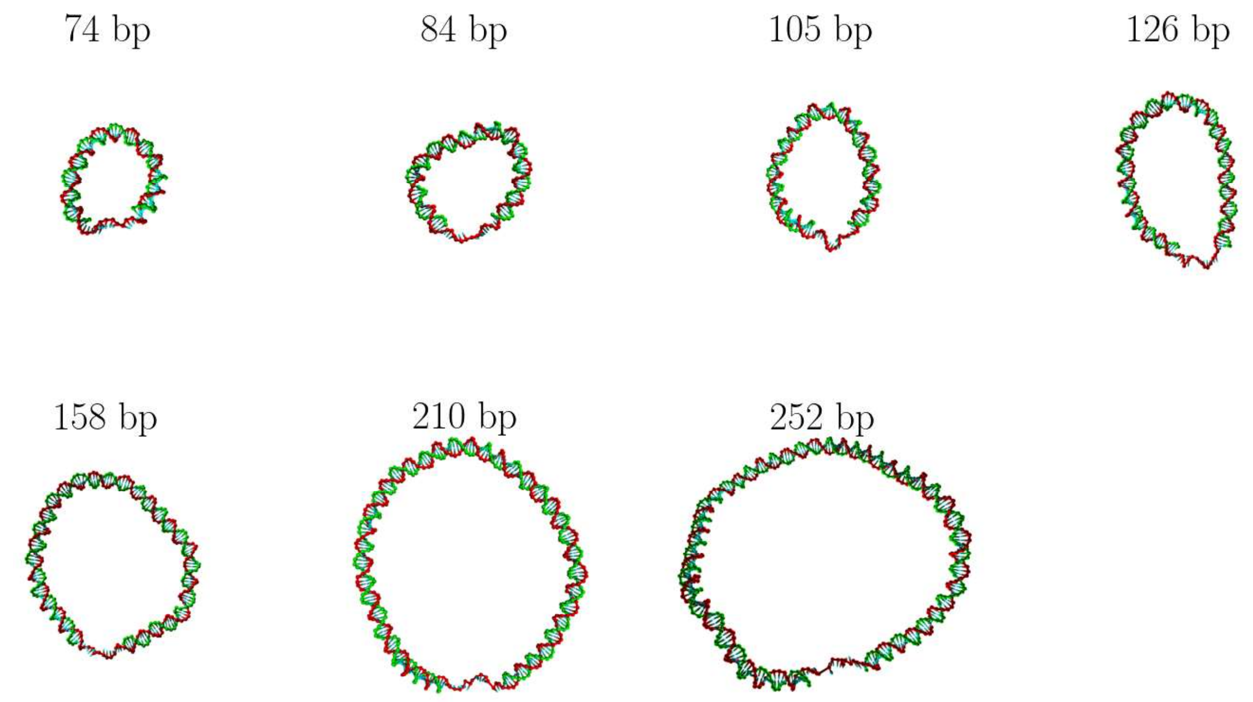


Supplementary Figure S3. Sample oxDNA2 configurations of all experimentally measured DNA bow sizes. Each configuration depicts the DNA bow in its unbound state. The label above each molecule specifies the length of the dsDNA bow arc (in base pairs). All constructs feature a 15 nt ssDNA strand containing a 9 nt region targeted by an 8-9 nt probe.

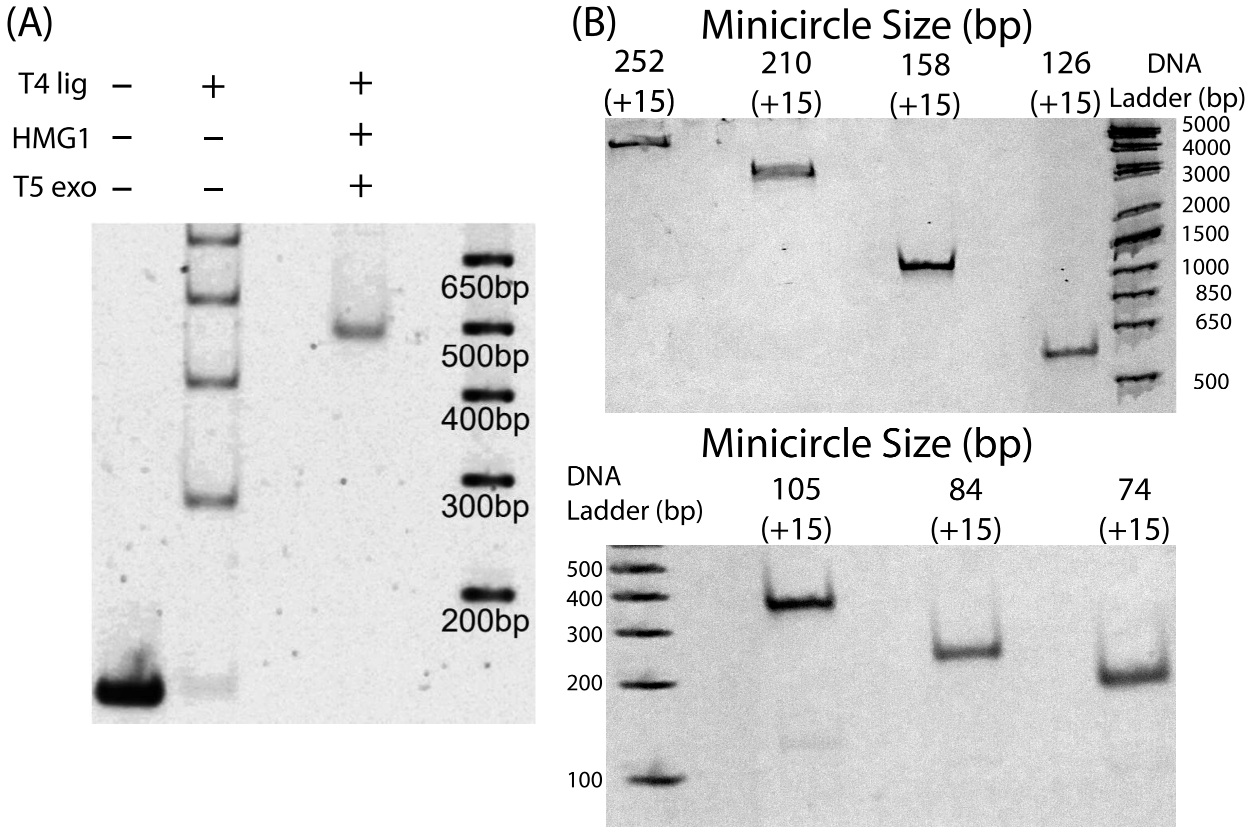


Supplementary Figure S4. **(A)** Linear DNA molecules with phosphorylated 5' ends were bent with bending protein HMG1 and self-ligated. T5 exonuclease was then added to digest unwanted polymer fragments. **(B)** The remaining circular DNA was then purified with ethanol precipitation and nicked on the unmodified strand with Nb.BbvCI. Nicked minicircle bands were analyzed and extracted using polyacrylamide gel electrophoresis (6%, 29:1 acrylamide to bis-acrylamide in 0.5x TBE buffer). Note that the total minicircle size includes the 15 bp target strand segment, which is not included in the bow arc length (74 bp to 252 bp. After each circle size was inspected via PAGE, DNA minicircles were extracted overnight via "crush-and-soak" and concentrated with ethanol precipitation.

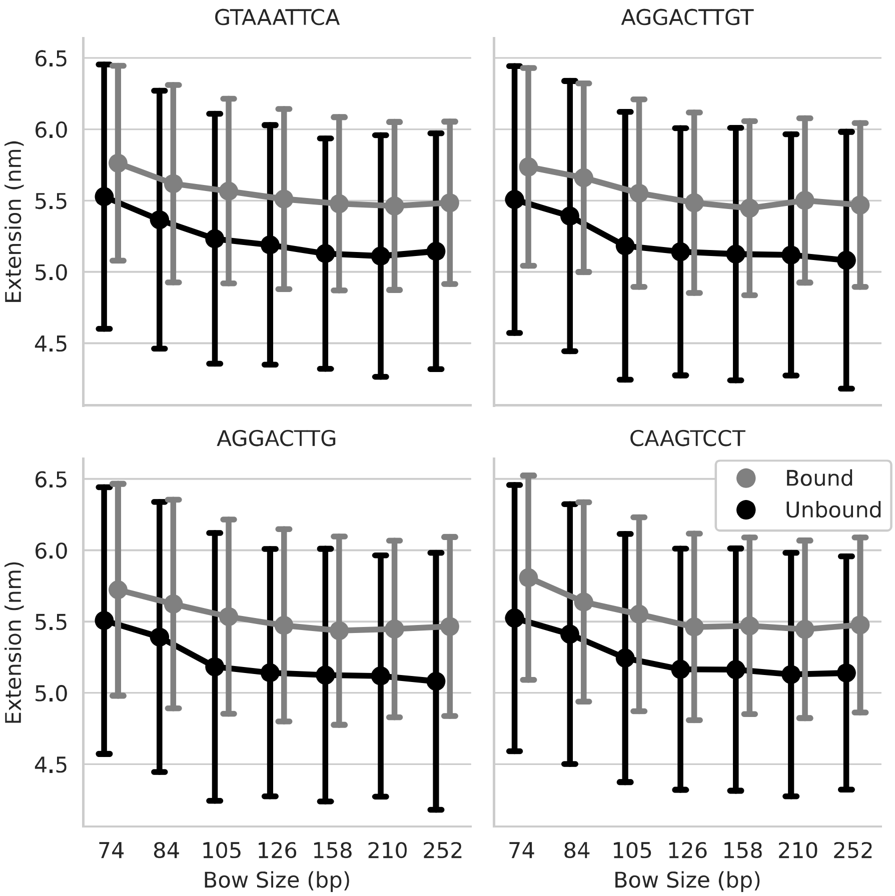


Supplementary Figure S5. Mean ($\overline{x}$) and standard deviation values ($\sigma(x)$) of the end-to-end distance $x$ for each DNA bow size, in both the probe-bound and probe-unbound states. Each MD simulation was performed for $7.5 \times 10^7$ steps with $dt = 15.2\,\text{fs}$, totaling $t = 1.14\,\mu\text{s}$. Extension values were measured every 1000 steps, collecting $n = 7.5 \times 10^4$ values in total.

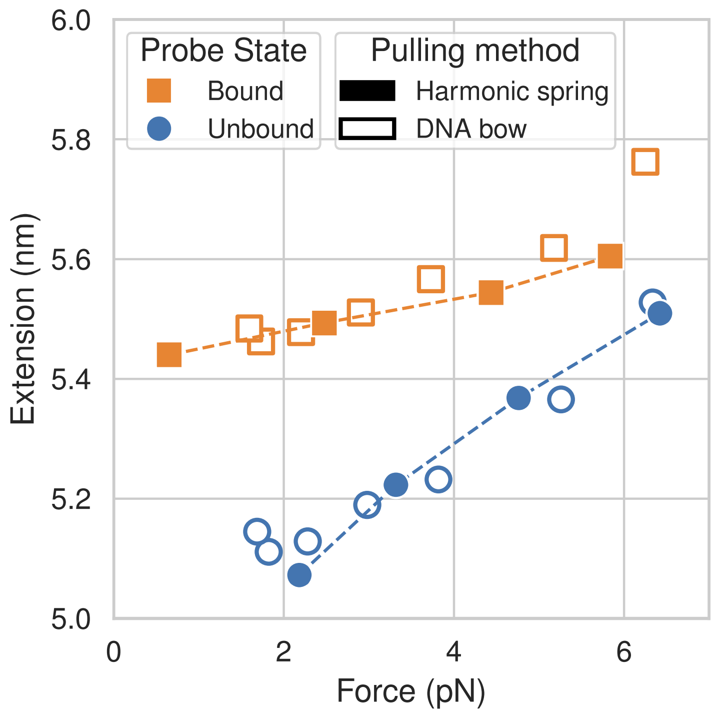


Supplementary Figure S6. Force-extension behavior of the 17 nt target sequence T1 in both its probe-bound and probe-unbound states, extended by either a harmonic spring (filled markers) or a DNA bow (open markers). For the harmonic spring pulling method, we used the "mutual trap" external force tool provided with oxDNA to connect the terminal bases of the target with a weak spring.

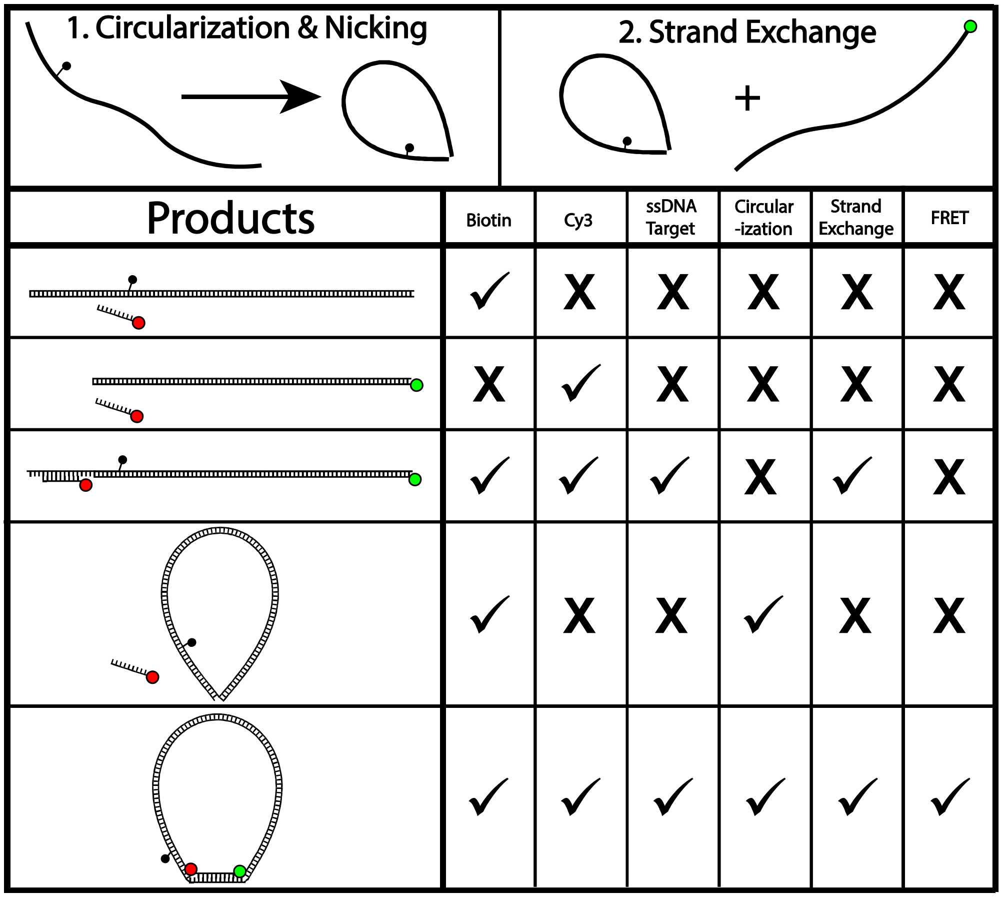


Supplementary Figure S7. Possible products created during bow construction. Nicked circular products were purified and mixed with Cy3-labeled linear molecules at a 1:4 ratio. The unmodified nicked strand is replaced via a strand exchange reaction which consists of heating the mixture to 95 °C and cooling to 4 °C gradually. By design, only the desired product (bottom row) is capable of generating a FRET signal. Incorrect purification of nicked circular strands (step one) would yield linear products during strand exchange; among these products, the target is either too far from the Cy3 dye to generate a FRET signal upon probe binding, or the target is absent entirely. Circular molecules that do not replace the unmodified strand during the strand exchange reaction (step two) are not donor-labeled, nor do they have an exposed acceptor-probe target, and therefore also cannot generate a FRET signal.

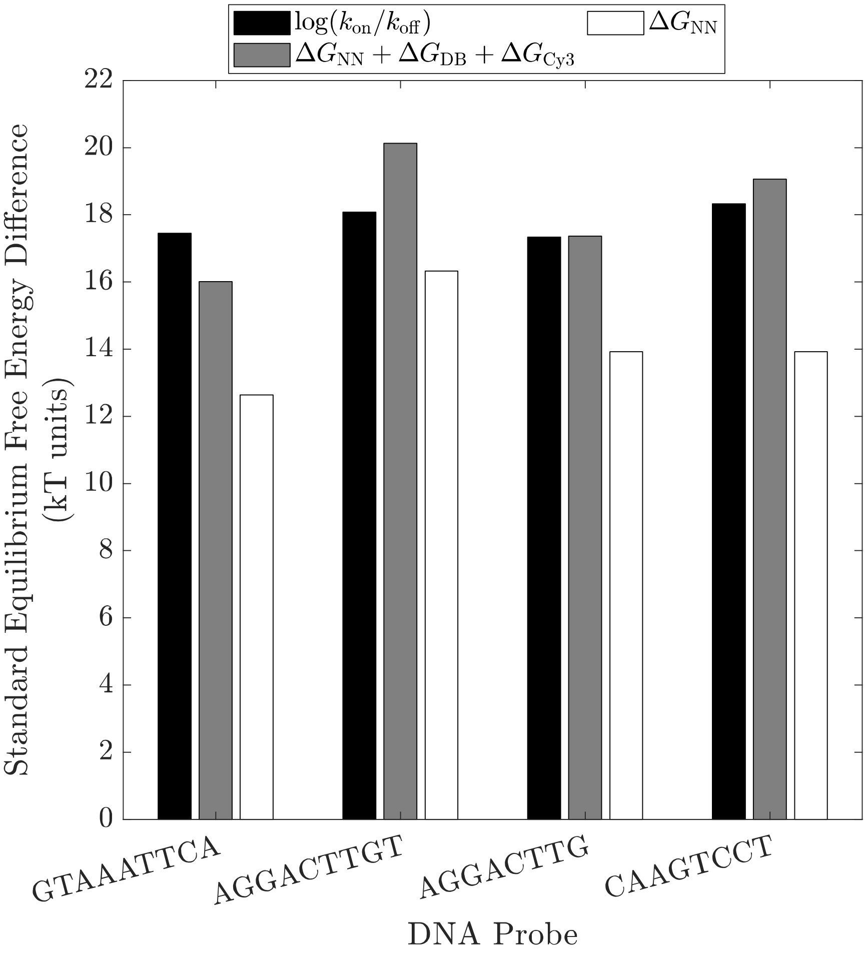


Supplementary Figure S8. A comparison of the measured standard equilibrium free energy difference values of all DNA probes to their corresponding nearest-neighbor predictions. The free energy difference, $\Delta G = \log(k_{on}/k_{off})$, was calculated using the average $k_{off}$ and $k_{on}$ values observed for the 252 bp DNA bow (black bars). The predicted free energy difference of a freely diffusing 8 bp to 9 bp duplex, $\Delta G_{NN}$, was then calculated using published nearest-neighbor thermodynamic parameters (white bars) [87]. We then modified this estimate to correct for our experimental conditions by adding the energy contribution of dangling base stacking interactions $\Delta G_{DB}$ [68] as well as the energy contribution of a Cy3 dye attachment $\Delta G_{Cy3}$ (gray bars) [72]. Both estimates were calculated at a temperature 22 °C to match our experiment. The resulting sum was corrected for the monovalent cation concentration of our buffer ($[Mono^+] = [Na^+] + [Tris^+] = 150$ mM) using the calibration formula published by SantaLucia Jr. Note that this estimate assumes that half of Tris molecules are protonated [88].

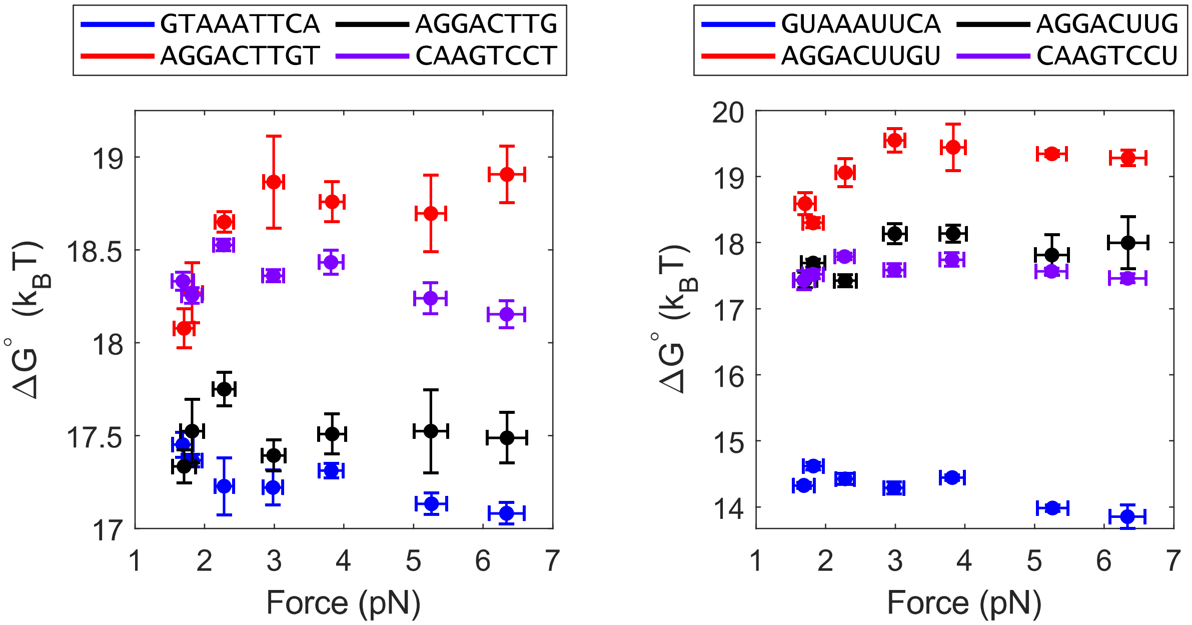


Supplementary Figure S9. Force dependence of the equilibrium free energy difference $\Delta G = \log(k_{\mathrm{on}}/k_{\mathrm{off}})$ between the bound and unbound states. Here, $\Delta G$ is defined as the additional free energy of the unbound state relative to the bound state. The force exerted by each bow was calculated with Equation 2, using the mean extension $\overline{x}$ of the bow's end-to-end distance distribution. Vertical error bars represent the standard error of the mean; horizontal error bars were calculated using $\frac{\partial f(x)}{\partial x}\Big|_{\overline{x}} \cdot \sigma(x)$, where $\overline{x}$ and $\sigma(x)$ are the mean and standard deviation of the bow's end-to-end distance distribution.

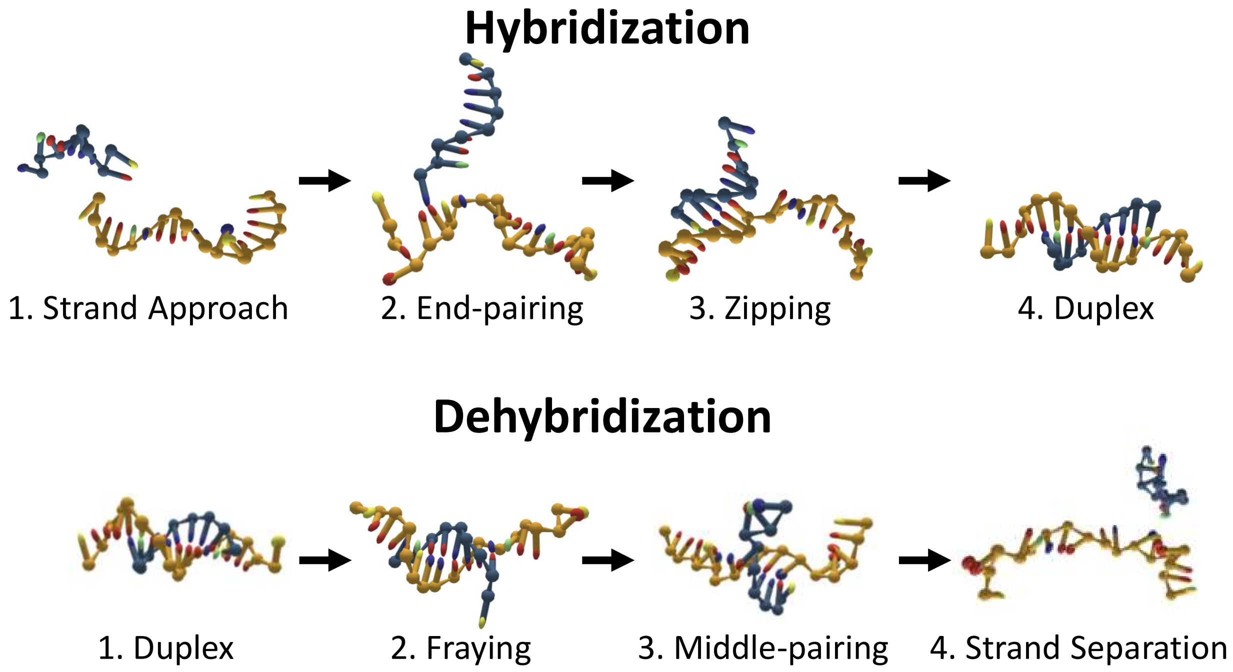


Supplementary Figure S10. Flowchart of hybridization and dehybridization reactions. During a typical binding transition, two strands approach one another and form an initial base pair at their terminal ends; afterwards, the two strands “zip” together in a linear fashion. During a typical unbinding transition, the base pairs at the ends of the two strands fray and separate, continuing inward until one last base pair remains near the center; after this final middle-pair melts, the strand separates.

| **DNA bow arc duplex segments (5′ to 3′)** | |
|---|---|
| 74 bp | GACTCCCCACTCGTCGTACGAGGTCGCACACGCCCCACACCCAGACCTCCCTGCCCTGGTACCTC<br>AGCACTGAG |
| 84 bp | GACTCCCCACTCGTCGTACGCAACGAGGTCGCACACGCCCCACACCCAGACCTCCCTGCGAGCGC<br>CTGGTACCTCAGCACTGAG |
| 105 bp | GACTCCCCACTCGTCGTACGATCGCCATGGCAACGAGGTCGCACACGCCCCACACCCAGACCTCC<br>CTGCGAGCGGGCATGGGTACCCTGGTACCTCAGCACTGAG |
| 126 bp | GACTCCCCACTCGTCGTACCACCCACGCGCGATCGCCATGGCAACGAGGTCGCACACGCCCCACA<br>CCCAGACCTCCCTGCGAGCGGGCATGGGTACAATGTCCCCGCCTGGTACCTCAGCACTGAG |
| 158 bp | GACTCCCCACTCGTCGTACGTTTGGGGAAAGACCACACCCACGCGCGATCGCCATGGCAACGAGG<br>TCGCACACGCCCCACACCCAGACCTCCCTGCGAGCGGGCATGGGTACAATGTCCCCGTTGCCACA<br>GAGACCACCCTGGTACCTCAGCACTGAG |
| 210 bp | GACTCCCCACTCGTCGTACTGCGAAATCCGGAGCAACGGGCAACCGTTTGGGGAAAGACCACACC<br>CACGCGCGATCGCCATGGCAACGAGGTCGCACACGCCCCACACCCAGACCTCCCTGCGAGCGGGC<br>ATGGGTACAATGTCCCCGTTGCCACAGAGACCACTTCGTAGCACAGCGCAGAGCGTAGCGCCTGG<br>TACCTCAGCACTGAG |
| 252 bp | GACTCCCCACTCGTCGTACTTTTTGTTTACGCGACAACTATGCGAAATCCGGAGCAACGGGCAAC<br>CGTTTGGGGAAAGACCACACCCACGCGCGATCGCCATGGCAACGAGGTCGCACACGCCCCACACC<br>CAGACCTCCCTGCGAGCGGGCATGGGTACAATGTCCCCGTTGCCACAGAGACCACTTCGTAGCAC<br>AGCGCAGAGCGTAGCGTGTTGTTGCTGCTGACAAAAGCCTGGTACCTCAGCACTGAG |
| **Primers for making DNA force assay duplex segments (5′ to 3′)** | |
| | ↓ 20 nt |
| 74 Forward | GACTCCCCACTCGTCGTACGAGGTCGCACACGCC |
| 84 Forward | GACTCCCCACTCGTCGTACGCAACGAGGTCGCACAC |
| 105 Forward | GACTCCCCACTCGTCGTACGATCGCCATGGCAACG |
| 126 Forward | GACTCCCCACTCGTCGTACCACCCACGCGCGAT |
| 158 Forward | GACTCCCCACTCGTCGTACGTTTGGGGAAAGACCACAC |
| 210 Forward | GACTCCCCACTCGTCGTACTGCGAAATCCGGAGCA |
| 252 Forward | GACTCCCCACTCGTCGTACTTTTTGTTTACGCGACAACTATG |
| 74 Reverse | CTCAGTGCTGAGGTACCAGGGCAGGGAGGTCTGGGTG |
| 84 Reverse | CTCAGTGCTGAGGTACCAGGCGCTCGCAGGGAGGT |
| 105 Reverse | CTCAGTGCTGAGGTACCAGGGTACCCATGCCCGCTC |
| 126 Reverse | CTCAGTGCTGAGGTACCAGGCGGGGACATTGTACCCATG |
| 158 Reverse | CTCAGTGCTGAGGTACCAGGGTGGTCTCTGTGGCAACG |
| 210 Reverse | CTCAGTGCTGAGGTACCAGGCGCTACGCTCTGCGCT |
| 252 Reverse | CTCAGTGCTGAGGTACCAGGCTTTTGTCAGCAGCAACAACA |
| **Primers for making circular molecules, target segment underlined (5′ to 3′)** | |
| T1 | [Phos]TTTTGAATTTACTTTGACTCCCCAC[BiotindT]CGTCGTAC |
| T2 & T3 | [Phos]TTTACAAGTCCTTTTGACTCCCCAC[BiotindT]CGTCGTAC |

<table>
<tr><td>T4</td><td>[Phos]TTTAGGACTTGTTTTGACTCCCCAC[BiotindT]CGTCGTAC</td></tr>
<tr><td>Reverse</td><td>[Phos]CTCAGTGC|TGAGGTACCAGG</td></tr>
<tr><td colspan="2">Primers for making Cy3-labeled molecules for strand exchange (5′ to 3′)</td></tr>
<tr><td>Forward</td><td>GACTCCCCACTCGTCGTAC</td></tr>
<tr><td>Reverse (Cy3)</td><td>[Cy3]CTCAGTGCTGAGGTACCAGG</td></tr>
<tr><td colspan="2">Cy5 acceptor probes (5′ to 3′)</td></tr>
<tr><td colspan="2">DNA and RNA probes for smFRET experiments (5′ to 3′)</td></tr>
<tr><td>P1-DNA</td><td>[Cy5]GTAAATTCA</td></tr>
<tr><td>P1-RNA</td><td>[Cy5]GUAAAUUCA</td></tr>
<tr><td>P2-DNA</td><td>[Cy5]AGGACTTGT</td></tr>
<tr><td>P2-RNA</td><td>[Cy5]AGGACUUGU</td></tr>
<tr><td>P3-DNA</td><td>[Cy5]AGGACTTG</td></tr>
<tr><td>P3-RNA</td><td>[Cy5]AGGACUUG</td></tr>
<tr><td>P4-DNA</td><td>[Cy5]CAAGTCCT</td></tr>
<tr><td>P4-RNA</td><td>[Cy5]CAAGUCCU</td></tr>
<tr><td colspan="2">FFS simulation sequences, (5′ to 3′)</td></tr>
<tr><td>P1-DNA</td><td>GTAAATTCA</td></tr>
<tr><td>T1</td><td>GTTTTGAATTTACTTTG</td></tr>
</table>

Supplementary Table S1: List of DNA sequences, PCR primers, and DNA/RNA probes. All bow arc duplex segments are sourced from yeast genomic DNA, and extended to include common adapter sequences on each end. Forward primers for making circular DNA include the 15 nt sequence containing the 9 nt ssDNA complementary target segment (underlined); the reverse primer for making circular DNA includes the nick site (marked with a vertical line "|"). DNA and RNA probes were added to imaging buffer at 20 nM during smFRET experiments to measure unbinding ($k_{\mathrm{off}}$) and binding rates ($k_{\mathrm{on}}$). Note that DNA target sequences used for FFS simulations include an additional nucleotide at each end, matching the letter of the terminal bases in the dsDNA portion of bow constructs.

| End-to-end extension (nm), $\overline{x} \pm \sigma(x)$ | | | | | | | |
|---|---|---|---|---|---|---|---|
| Unbound state | | | | | | | |
| Target | 74 bp | 84 bp | 105 bp | 126 bp | 158 bp | 210 bp | 252 bp |
| 1 | $5.5 \pm 0.9$ | $5.4 \pm 0.9$ | $5.2 \pm 0.9$ | $5.2 \pm 0.8$ | $5.1 \pm 0.8$ | $5.1 \pm 0.8$ | $5.1 \pm 0.8$ |
| 2 & 3 | $5.5 \pm 0.9$ | $5.4 \pm 0.9$ | $5.2 \pm 0.9$ | $5.1 \pm 0.9$ | $5.1 \pm 0.9$ | $5.1 \pm 0.8$ | $5.1 \pm 0.9$ |
| 4 | $5.5 \pm 0.9$ | $5.4 \pm 0.9$ | $5.2 \pm 0.9$ | $5.2 \pm 0.8$ | $5.2 \pm 0.8$ | $5.1 \pm 0.9$ | $5.1 \pm 0.8$ |
| Bound state | | | | | | | |
| Target | 74 bp | 84 bp | 105 bp | 126 bp | 158 bp | 210 bp | 252 bp |
| 1 | $5.8 \pm 0.7$ | $5.6 \pm 0.7$ | $5.6 \pm 0.6$ | $5.5 \pm 0.6$ | $5.5 \pm 0.6$ | $5.5 \pm 0.6$ | $5.5 \pm 0.6$ |
| 2 | $5.7 \pm 0.7$ | $5.7 \pm 0.7$ | $5.6 \pm 0.7$ | $5.5 \pm 0.6$ | $5.4 \pm 0.6$ | $5.5 \pm 0.6$ | $5.5 \pm 0.6$ |
| 3 | $5.7 \pm 0.7$ | $5.6 \pm 0.7$ | $5.5 \pm 0.7$ | $5.5 \pm 0.7$ | $5.4 \pm 0.7$ | $5.4 \pm 0.6$ | $5.5 \pm 0.6$ |
| 4 | $5.8 \pm 0.7$ | $5.6 \pm 0.7$ | $5.6 \pm 0.7$ | $5.5 \pm 0.7$ | $5.5 \pm 0.6$ | $5.4 \pm 0.6$ | $5.5 \pm 0.6$ |

Supplementary Table S2. Mean ($\overline{x}$) and standard deviation values ($\sigma(x)$) of the end-to-end distance $x$ of each DNA bow in both the probe-bound and probe-unbound state. Values were calculated using the measured $x$ distance values of $7.5 \times 10^4$ configurations saved over $t = 1.14\,\mu s$ of simulation time. $x$ is defined as the distance between backbone sites on the terminal bases of the elastic arc that are covalently linked to the ssDNA target segment.

| Order Parameter Q | Number of base pairs $n$ ($E < E_0$) |
|---|---|
| $Q = -1$ | $n = 9$ |
| $Q = 0$ | $n = 8$ |
| $Q = 1$ | $n = 7$ |
| $Q = 2$ | $n = 6$ |
| $Q = 3$ | $n = 5$ |
| $Q = 4$ | $n = 4$ |
| $Q = 5$ | $n = 3$ |
| $Q = 6$ | $n = 2$ |
| $Q = 7$ | $n = 1$ |
| $Q = 8$ | $n = 0$ |

Supplementary Table S3. Order parameter definitions for FFS unbinding simulations. Each interface is defined by the value of a corresponding order parameter (e.g. separation $d$ or base pairs). Base pairs are defined as when two complementary nucleotides have a hydrogen bonding energy lower than the energy scale $E_0 = -0.6\,\text{kcal/mol}$. Note that base pairs formed in misaligned duplex structures are not counted.

| Order Parameter Q | Minimum Separation $d$/nm | Base pairs $n$ ($E < E_0$) |
|---|---|---|
| $Q = -2$ | $d > 3.41$ | - |
| $Q = -1$ | $3.41 \geq d > 1.70$ | - |
| $Q = 0$ | $1.70 \geq d > 0.85$ | - |
| $Q = 1$ | $d \leq 0.85$ | $n = 0$ |
| $Q = 2$ | - | $n = 1$ |
| $Q = 3$ | - | $n = 2$ |
| $Q = 4$ | - | $n = 3$ |
| $Q = 5$ | - | $n = 4$ |
| $Q = 6$ | - | $n = 5$ |
| $Q = 7$ | - | $n = 6$ |
| $Q = 8$ | - | $n = 7$ |
| $Q = 9$ | - | $n = 8$ |
| $Q = 10$ | - | $n = 9$ |

Supplementary Table S4. Order parameter definitions for FFS binding simulation. Each interface of the FFS simulation is defined using an order parameter value (e.g. separation $d$ or base pairs $n$). The symbol '-' indicates that there is no constraint on the particular coordinate for the given interface. Minimum separation is defined as the minimum distance between any two complementary bases on the target and probe strands. Base pairs are defined as when two complementary nucleotides have a hydrogen bonding energy lower than the energy scale $E_0 = -0.6\,\text{kcal/mol}$. Note that misaligned duplex structures are not counted as base pairs.

| **FFS results for probe P1 (GTAAATTCA)** | | | | | | | | | |
|---|---|---|---|---|---|---|---|---|---|
| | | **Reaction type** | | | | | | | |
| | | Unbinding | | | | Binding | | | |
| | | **Target strand end-to-end fixed extension value** | | | | | | | |
| | | 5.5 nm | | | | 5.1 nm | | | |
| **Trial** | **Goal Interface** | **Number of forward crossings** | | | | | | | |
| 1 | $\lambda_0$ | 30017 | | | | 30002 | | | |
| 2 | | 60013 | | | | 60001 | | | |
| 3 | | 60015 | | | | 60000 | | | |
| **Trial** | **Goal Interface** | **Initial Flux** ($ns^{-1}$) | | | | | | | |
| 1 | $\lambda_0$ | 57.3 | | | | 0.515 | | | |
| 2 | | 56.6 | | | | 0.518 | | | |
| 3 | | 57.4 | | | | 0.520 | | | |
| **Trial** | **Goal Interface** | **Successes** | **Prune Successes** | **Attempts** | **Prob.** | **Successes** | **Prune Successes** | **Attempts** | **Prob.** |
| 1 | $\lambda_1$ | 30001 | - | 857297 | 0.035 | 30001 | - | 1309017 | 0.023 |
| 2 | | 30000 | - | 861360 | 0.035 | 30000 | - | 1308931 | 0.023 |
| 3 | | 30001 | - | 850404 | 0.035 | 30000 | - | 1302569 | 0.023 |
| 1 | $\lambda_2$ | 30002 | 15223 | 1299683 | 0.058 | 15585 | 859 | 6864816 | 0.003 |
| 2 | | 30001 | 15048 | 1284311 | 0.059 | 12230 | 928 | 5173098 | 0.003 |
| 3 | | 30000 | 15167 | 1276431 | 0.059 | 13688 | 924 | 5733836 | 0.003 |
| 1 | $\lambda_3$ | 30004 | 17169 | 1543954 | 0.053 | 30001 | 93 | 174849 | 0.173 |
| 2 | | 30003 | 17306 | 1539043 | 0.053 | 30001 | 113 | 168188 | 0.18 |
| 3 | | 30001 | 17395 | 1491069 | 0.055 | 30000 | 112 | 165547 | 0.183 |
| 1 | $\lambda_4$ | 30001 | 17373 | 1837336 | 0.045 | 30000 | 17857 | 135201 | 0.618 |
| 2 | | 30002 | 17244 | 1799833 | 0.045 | 30001 | 17697 | 130722 | 0.636 |
| 3 | | 30003 | 17451 | 1728580 | 0.048 | 30002 | 17233 | 125808 | 0.649 |
| 1 | $\lambda_5$ | 30001 | 17501 | 1930017 | 0.043 | 30012 | 17036 | 194725 | 0.942 |
| 2 | | 30001 | 17707 | 1871701 | 0.044 | 30005 | 16393 | 185757 | 0.956 |
| 3 | | 30000 | 18032 | 1829295 | 0.046 | 30012 | 15795 | 180810 | 0.952 |
| 1 | $\lambda_6$ | 30000 | 19898 | 1693959 | 0.053 | 30018 | 15787 | 173298 | 0.993 |
| 2 | | 30000 | 20882 | 1489686 | 0.062 | 30006 | 16457 | 179119 | 0.994 |
| 3 | | 30001 | 20541 | 1565933 | 0.059 | 30014 | 16098 | 176816 | 0.989 |
| 1 | $\lambda_7$ | 10011 | 6924 | 554063 | 0.056 | 30021 | 22227 | 96261 | 1.005 |
| 2 | | 10004 | 7206 | 499387 | 0.063 | 30010 | 22622 | 98389 | 0.995 |
| 3 | | 6796 | 4788 | 370763 | 0.057 | 30022 | 22178 | 97357 | 0.992 |
| 1 | $\lambda_8$ | 10001 | 7386 | 832578 | 0.039 | 30022 | 20175 | 90038 | 1.006 |
| 2 | | 9114 | 6605 | 811829 | 0.036 | 30011 | 20179 | 90944 | 0.996 |
| 3 | | 5000 | 3596 | 382574 | 0.041 | 30021 | 20319 | 91490 | 0.994 |
| 1 | $\lambda_9$ | 10007 | 5000 | 256882 | 0.097 | 30022 | 18749 | 86244 | 1.0 |
| 2 | | 10001 | 5030 | 255935 | 0.098 | 30011 | 18767 | 86104 | 1.002 |
| 3 | | 5002 | 2576 | 127468 | 0.1 | 30022 | 18803 | 86856 | 0.995 |
| 1 | $\lambda_{10}$ | - | - | - | - | 30022 | 18571 | 85373 | 1.004 |
| 2 | | - | - | - | - | 30023 | 18355 | 85408 | 0.996 |
| 3 | | - | - | - | - | 30022 | 18628 | 85253 | 1.008 |

| $k_{AB}$ | $9.9\,\mathrm{min}^{-1}$ | $2.3\,\mathrm{s}^{-1}\,\mu\mathrm{M}^{-1}$ |
|---|---|---|

Supplementary Table S5. Forward flux simulation results for binding and unbinding reactions. The "Successes" column specifies the number of configurations that successfully cross the goal interface $\lambda_Q$ coming from $\lambda_{Q-1}$; "Prune Successes" specifies the number of configurations that survive pruning upon traveling backward to $\lambda_{Q-2}$ from $\lambda_{Q-1}$, and afterward cross $\lambda_Q$. Trajectories that revert to $\lambda_{Q-2}$ were pruned with a $p = 0.75$ probability. "Attempts" specifies the total number of trajectories started at $\lambda_{Q-1}$. The probability ("Prob") of crossing $\lambda_0$ was calculated using Equation $S5$, while the probability of crossing the remaining interfaces was calculated using Equation $S6$. The final rate $k_{AB}$ was then calculated using Equation 3.
